# Recurrent gene flow events shaped the diversification of the clownfish skunk complex

**DOI:** 10.1101/2023.10.24.562491

**Authors:** Anna Marcionetti, Joris A. M. Bertrand, Fabio Cortesi, Giulia F. A. Donati, Sara Heim, Filip Huyghe, Marc Kochzius, Loïc Pellissier, Nicolas Salamin

**Author notes:** co-first authorship (Anna Marcionetti, Joris A. M. Bertrand). Author for Correspondence: Nicolas Salamin, Department of Computational Biology, University of Lausanne, 1015 Lausanne, Switzerland.

## Abstract

Clownfish (subfamily Amphiprioninae) are an iconic group of coral reef fish that evolved a mutualistic interaction with sea anemones, which was shown to have triggered the adaptive radiation of the group. Within clownfishes, the skunk complex is particularly interesting as, besides ecological speciation, gene flow between species and hybrid speciation are suggested to have shaped the diversification of the group. We investigated, for the first time, the mechanisms underlying the diversification of this complex. By taking advantage of their disjunct geographical distribution, we obtained whole-genome data of sympatric and allopatric populations of the three main species of the complex (*Amphiprion akallopisos*, *A. perideraion* and *A. sandaracinos*). We examined the population structure, genomic divergence patterns and introgression signals, and performed demographic modeling to identify the most realistic diversification scenario. We excluded scenarios of strict isolation, of hybrid origin of *A. sandaracinos*, and ruled out the presence of extensive gene flow in sympatry. We discovered moderate gene flow from *A. perideraion* to the ancestor of *A. akallopisos + A. sandaracinos* and weak gene flow between the species in the Indo-Australian Archipelago throughout the diversification process of the group. We identified introgressed regions in *A. sandaracinos* and detected two large regions of high divergence in *A. perideraion*, likely maintained by the disruption of recombination. Altogether, our results show that ancestral hybridization events shaped the group’s diversification. However, more recent gene flow is less pervasive than initially thought and suggests a role of host repartition or behavioral barriers in maintaining the genetic identity of the species in sympatry.

## INTRODUCTION

The underlying causes and mechanisms of organisms’ adaptation, diversification, and the emergence of new species, have been fundamental questions in evolutionary biology (Losos et al., 2013). Answering these questions is not straightforward, because the speciation process is complex and involves different and interplaying mechanisms, such as geographic isolation, genetic drift, natural selection, and sexual selection (Gavrilets, 2004; Coyne, 2004; Gavrilets & Losos, 2009; Schluter, 2009; Nosil et al., 2009). Since access to genomic data is increasing rapidly, the investigation of mechanisms underlying diversification and speciation has been extended to numerous taxa, environments, and evolutionary stages (Seehausen et al., 2014; Berner & Salzburger, 2015; Cerca et al., 2023). Studies have identified loci promoting ecological divergence and, eventually, speciation (e.g., Malinsky et al., 2015; Payseur & Rieseberg, 2016). Likewise, the prevalence and importance of hybridization in the evolution of species has become evident (Abbott et al., 2013; Payseur & Rieseberg, 2016; Taylor & Larson, 2019). While hybridization was traditionally seen as a process limiting diversification (Mayr, 1963), it is now recognized to promote ecological adaptation and speciation (Abbott et al., 2013; Taylor & Larson, 2019) and it is thus, a pervasive key element of adaptive radiations (Berner & Salzburger, 2015; Cerca et al., 2023). Hybrid speciation (Olave et al., 2022) and ancient hybridization events that fueled diversification (Meier et al., 2017; Svardal et al., 2020; Meier et al., 2023) have been observed in cichlid fishes, while introgressive hybridization linked to adaptive divergence has been found in *Heliconius* butterflies (Nadeau et al., 2013).

These genomic studies in cichlids and butterflies have contributed considerably to our understanding of the molecular bases for diversification and speciation (Seehausen et al., 2014; Berner & Salzburger, 2015). However, they have also highlighted that more comparative studies considering different organisms at various stages of divergence are needed to fully grasp the interplay between genomic properties, ecology, geography, and demographic history in shaping species diversification (Seehausen et al., 2014; Wolf & Ellegren, 2017; Campbell et al., 2018). The skunk complex (or *akallopisos* group) within the clownfish genus *Amphiprion* (family Pomacentridae) lends itself to this purpose. This complex is composed of three main species distributed across the Indo-Pacific Ocean, *Amphiprion akallopisos*, *A. sandaracinos* and *A. perideraion* (Fig. 1A; Fautin & Allen., 1997), as well as a fourth species that is endemic to Fiji, Tonga, Samoa, and Wallis Island (*A. pacificus*; Allen et al., 2010). Like all clownfishes, they maintain a mutualistic interaction with sea anemones (Fautin & Allen., 1997). However, the members of the skunk complex show divergence in host use. *A. perideraion* is a generalist that can interact with four sea anemone species, while *A. sandaracinos* and *A. akallopisos* only associate with two host species (Supplementary Table S1; Fautin & Allen, 1997; Litsios et al., 2012). The group diversified in the Indo-Australian Archipelago (IAA) ca. 1 to 5 MYA (Cowman et al., 2013; Litsios et al., 2012; Frédérich et al., 2013; Litsios et al., 2014; Rabosky et al., 2018). Today, the skunk species have a vast and only-partially overlapping geographical distribution (Fig. 1A). Although the three species still co-occur in the IAA, *A. akallopisos* has a disjunct distribution and can also be found in the Western Indian Ocean (WIO), while *A. perideraion* is observed further east, reaching the New Caledonian (NC) archipelago (Allen, 1991; Fig. 1A). Based on cytonuclear inconsistencies detected between mitochondrial and nuclear phylogenetic trees, ancestral hybridization events are thought to have shaped the diversification of the complex, and a hybrid origin has been suggested for *A. sandaracinos* (Litsios & Salamin, 2014). Furthermore, hybrids between species of the skunk complex are commonly observed in the aquarium trade. Contemporary gene flow between the three species in the IAA is also likely, as suggested by the low interspecific barcode variation for the species in this region (Steinke et al., 2009). However, whole genome characterization is required to test the hypothesis that hybridization played a major role in the evolution of the skunk complex and the origin of its species.

**Figure 1.**
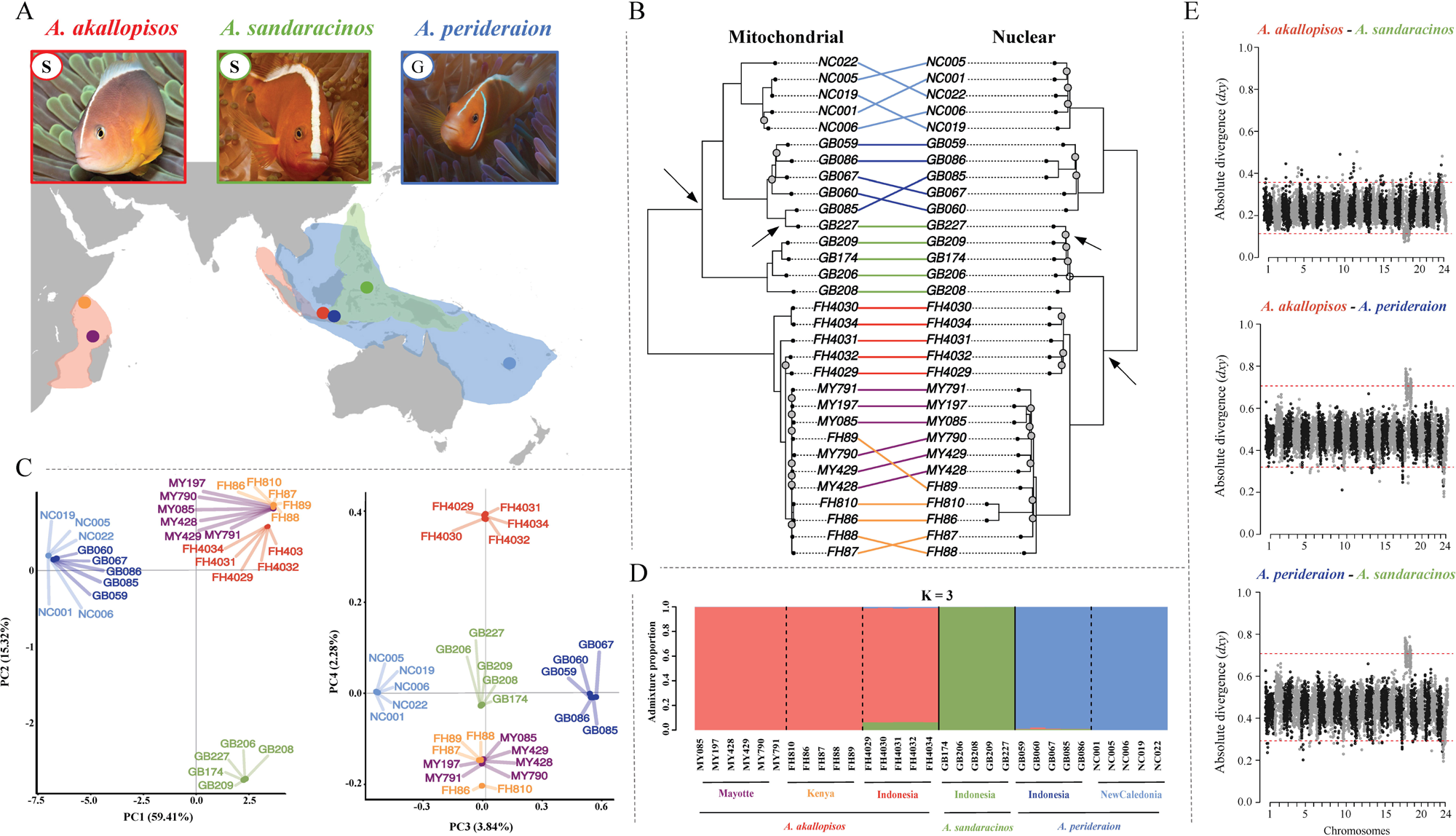
The three *Amphiprion* species considered in this study, their geographical distributions, and the locations of the sampled populations (**A**). The classification based on the host sea anemone usage of the three species is reported, with S=Specialist and G=Generalist (Supplementary Table S1). Cytonuclear discordance is observed (**B)**, but the species and populations are overall well differentiated, as shown by the PCA (**C**) and the admixture plot (**D**). The *d_xy_* along the 24 chromosomes for the different species pairs also show a clear genetic divergence of the populations, but two large outlier regions are observed on chromosome 18 (**E**). *A. akallopisos*, *A. sandaracinos* and *A. perideraion* samples and populations are reported in shades of red, green, and blue, respectively. **(B)** The mitochondrial phylogenetic tree was obtained with RAxML from the alignment of the whole mitochondrial genomes (16,747 bp), and the nuclear tree was obtained with RAxML and ASTRAL-III from 29,793,603 SNPs. Grey dots indicate bootstrap support lower than 0.8, and the arrows pinpoint the cytonuclear inconsistencies. The phylogenetic trees were rooted with *A. percula* as outgroup, which was removed from the plot. (**C, D**) The PCA and admixture results were obtained with PCAngsd on 29,793,603 SNPs. In the PCA, the first two components separate samples according to the species, whereas PC3 and PC4 separate the populations of *A. perideraion* and *A. akallopisos* according to geography. For the admixture analysis, multiple runs with different seeds for each K were performed to ensure convergence, and the highest likelihood solution was kept. The admixture plot represents the individuals ancestry proportions for the best number of ancestral populations K selected by PCAngsd (K=3). Plots for different K are available in Supplementary Fig. S3. (**E**) The absolute genetic divergence (*d_xy_*) along the 24 chromosomes for each species pair is reported for sliding windows of 3,000 variable sites. Plots for the Indonesian populations of each species are shown, but similar results were obtained for all populations (Supplementary Fig. S4). Red dotted lines represent the upper and lower 1% of the *d_xy_* distributions. Outlier regions of increased (*A. perideraion – A. akallopisos* and *A. perideraion – A. sandaracinos*) or decreased (*A. akallopisos – A. sandaracinos*) divergence are observed on chromosome 18.

In this study, we investigated the mechanisms driving the diversification of the *akallopisos* group by sampling and sequencing sympatric and allopatric populations of the three species. Our goal was to test whether *A. sandaracinos* originated through hybrid speciation or whether its diversification was characterized by ancestral hybridization events, as suggested by the cytonuclear discordance. Under the hypotheses of ancestral gene flow, ecological divergence in host usage (i.e., specialists (S) vs generalists (G), Fig. 1A) may have facilitated the speciation process. Ecological divergence in terms of the number of possible mutualistic sea anemone hosts is thought to be the driver for the ecological speciation and adaptive radiation in clownfishes, especially for the species with overlapping distributions centered in the IAA (Litsios et al., 2012). Additionally, we examined the occurrence of recent interspecific gene flow in the IAA. The presence of gene flow between populations of different species could result in the homogenization of their genomic divergence, which, in turn, would allow the identification of genomic regions of increased differentiation potentially involved in the ecological divergence or speciation of the complex (e.g., Nadeau et al., 2012; Bourgeois et al., 2020; Luzuriaga-Aveiga et al., 2021; Barrera-Guzmán et al., 2022). This study, therefore, offers a better understanding of the role of hybridization (ancestral and/or recent) and ecological divergence in the diversification of the skunk complex and, more broadly, in the radiation of clownfishes.

## MATERIAL AND METHODS

### Species selection and sampling

We obtained the overall geographic distribution of the species of the skunk complex from Allen (1991) and GBIF.org (GBIF Occurrence Download https://doi.org/10.15468/dl.rzpkl4, accessed in January 2016; Fig. 1A, Supplementary Fig. S1). We considered three species: *Amphiprion akallopisos, A. perideraion* and *A. sandaracinos*. We excluded the fourth one (*A. pacificus*) from the study because the geographic distribution of this species as of the most recent update in 2010, (Allen et al., 2010) is restricted to the vicinity of a handful of Pacific islands: Fidji, Tonga, Wallis, and Samoa. The three studied species show slight differences in color patterns (Fautin & Allen, 1997). *A. akallopisos* and *A. perideraion* have a white caudal fin and an orange pinkish body color, whereas the caudal fin and the body coloration of *A. sandaracinos* are orange (Fig. 1A). *A. sandaracinos* has a slightly longer white stripe than the other two species, while *A. perideraion* has an additional white bar on the head (Fig. 1A). *A. akallopisos* and *A. perideraion* have similarly shaped teeth (incisiform), which differ in *A. sandaracinos* (conical), potentially indicating slightly different ecological adaptations (Timm et al., 2008).

We collected individuals from the IAA region for each species. For *A. perideraion* and *A. akallopisos*, we also sampled populations at the margin of their geographic distribution, resulting in a total of three populations of *A. akallopisos* (Indonesia, Mayotte, and Kenya), two populations of *A. perideraion* (Indonesia and New Caledonia) and one population of *A. sandaracinos* (Indonesia). We sampled five to six individuals per population (Fig 1, Supplementary Table S2). Fish were caught in their host anemone using nets while Scuba diving. A piece of the caudal fin was taken and preserved in 96% ethanol immediately after the dive, in agreement with the permits delivered by the official institutions of the different countries. A fin clip of one *A. percula* individual from Lizard Island, Northern Great Barrier Reef, Australia, was obtained from Fabio Cortesi (Queensland Brain Institute, University of Queensland, Australia) and used as an outgroup for subsequent analyses.

### DNA extraction, library preparation and DNA sequencing

Genomic DNA (gDNA) was extracted from the sampled tissue using a DNeasy Blood and Tissue Kit (Qiagen, Hilden, Germany) following the manufacturer’s instructions. We prepared short-insert (350 bp) paired-end (PE) libraries from 100 ng of gDNA using a TruSeq Nano DNA LT Library Preparation Kit (Illumina) following the manufacturer’s instructions. The libraries corresponding to individuals of *A. sandaracinos*, *A. perideraion* from Indonesia, and the *A. percula* individual were sequenced on an Illumina HiSeq4000 at the Genomics Platform iGE3 (University of Geneva, Geneva), with a multiplex level of 8 individuals per lane (read length: 100 bp). The remaining libraries were sequenced on an Illumina HiSeq2000 at the Lausanne Genomic Technologies Facility, multiplexing five individuals per lane (read length: 100 bp).

### Reads processing, mapping, and variant calling

We removed adapter contamination from the raw reads with Cutadapt (v.1.13; Martin, 2011), trimmed the reads with Sickle (parameters: --qual-threshold 20, --length-threshold 40; v.1.33; Joshi & Fass, 2011) and verified their quality with FASTQC (v.0.11.5; Andrews, 2010). We mapped reads to the *A. percula* reference genome (INSDC Assembly <u>GCA_003047355.1</u>; Lehmann et al., 2018) using BWA (v.0.7.15; Li & Durbin, 2009) and generated mapping statistics with bamtools (command *stats*, v.2.4.1; Barnett et al., 2011) and Picard tools (command *CollectInsertSizeMetrics* v.2.4.1; http://broadinstitute.github.io/picard). We filtered the mapped reads with samtools (command *view*, parameters: -f 2 -F 256 -q 30; v.1.3; Li et al., 2009) to keep only primary alignments and proper pairs with a mapping quality higher than 30. We used *mergeReads* from ATLAS (v.0.9; Link et al., 2017) to remove redundant sequencing data originating from the overlap of paired reads and verified the absence of soft-clipped positions with *assessSoftClipping* from ATLAS (v.0.9; Link et al., 2017).

SNPs were called with the ATLAS pipeline (v.0.9; Link et al., 2017), which performs well at low-to-medium coverage and maintains a high accuracy in variant calling for moderately divergent species (Duchen & Salamin, 2020). Genotype likelihoods (GL) were computed with the *GLF* task (windows size of 0.1 Mb), and SNPs were subsequently called with the *majorMinor* task (MLE method). Window sizes of 0.1 Mb allow reducing the computational effort while maintaining GL accuracy when the average coverage is at least 4 (Kousathanas et al., 2016). We filtered out invariant sites, low-quality SNPs and singletons using vcftools (parameters: --minQ 40 --minDP 2 --max-missing 0.9 --max-meanDP 40 -- maxDP 40 --maf 0.02; v.0.1.15; Danecek et al., 2011). With vcftools, we also generated a second SNPs dataset by removing the *A. percula* reads and subsequently filtering out resulting monomorphic positions and singletons. We repeated the SNP calling as described above but excluding the outgroup to assess its potential influence on SNPs specific to the skunk complex. Results were consistent, with a difference of only 326,456 SNPs (2% of the total number of SNPs). We computed SNPs densities for different window sizes (100kb, 150kb, 200kb and 250kb) with vcftools (v.0.1.15; Danecek et al., 2011). Plotting was performed with the R package ggplot2 (v.3.0.0; Wickham, 2016).

### Mitochondrial genome assembly

Mitochondrial genomes were reconstructed with MITObim (v.1.9; Hahn et al., 2013) by either using barcode sequences to initiate the assembly or previously published mitochondrial genomes as reference. The sequences GB KJ833753 (mitochondrial genome; Li et al., 2015) and GB FJ582806 (COI gene; Steinke et al., 2009) were used for the mitochondrial genome reconstruction of *A. perideraion* samples. The sequences GB JF434730 (mitochondrial genome; Hubert et al., 2012) and JF434730 (COI gene; Hubert et al., 2012) were used for *A. akallopisos* and *A. perideraion* samples. We confirmed the consistency of the two reconstruction methods with Geneious (v.10.2.2; Kearse et al., 2012), and we manually inferred the circularity of the sequence.

### Phylogenetic reconstruction

The mitochondrial genome of one *A. percula* individual was obtained from NCBI (GB NC_023966.1; Tao et al., 2016). The mitochondrial genomes of all species were aligned using MAFFT (default parameters; v.7.450; Katoh & Standley, 2013). We visually checked the alignment to remove poorly aligned regions and reconstructed the mitochondrial phylogenetic tree with RAxML (GTR+Γ model, 100 bootstrap; v.8.2.12; Stamatakis, 2014).

For the nuclear dataset, we generated SNPs alignments using the script parseVCF (https://github.com/simonhmartin/genomics_general) and reconstructed phylogenetic trees for each chromosome separately using RAxML (GTR+Γ model, 100 bootstrap replicates, Lewis’ ascertainment bias correction; v.8.2.12; Stamatakis, 2014). The nuclear phylogenetic tree was then inferred with ASTRAL-III (v.5.6.3; Zhang et al., 2018) from the chromosomal phylogenetic trees. The two resulting trees (mitochondrial and nuclear) were plotted with the *cophylo* command of the R package phytools (v.0.6.44; Revell, 2012). The *A. percula* individual was used to root the two phylogenetic trees and was removed for further analyses. <u>Principal Component Analysis and Admixture</u>

We performed a Principal Component Analysis (PCA) and admixture analysis with PCAngsd (v.0.98; Meisner & Albrechtsen, 2018). The PCA was performed on the whole dataset and on each chromosome separately to check for disparities. In the admixture analysis, we considered two to five ancestral populations (K) and estimated individual ancestry coefficients by setting the parameter -e to the corresponding K-1 value, as recommended in the PCAngsd guidelines. For each K, the analysis was run five times with different seeds to ensure convergence, and the outcome with the highest likelihood was retained. The best K was automatically selected by PCAngsd based on the MAP test.

### Estimation of population genomic metrics

We investigated the overall genomic differentiation between the populations and species by estimating the average *F_st_* for each pair of populations using vcftools (v.0.1.15; Danecek et al., 2011; Supplementary Information S1). Sliding windows measures of relative (*F_st_*) and absolute (*d_xy_*) genetic divergence and nucleotide diversity (π) were calculated using *popgenWindows.py* (https://github.com/simonhmartin/genomics_general). To account for the variability in SNPs density along the genome, window sizes were defined based on the number of variable sites (parameter: --windType sites). We performed analyses with windows of 500 to 5,000 variable sites, corresponding to an average window size of approximately 25 to 250 kb. The results for different window sizes were consistent (Supplementary Table S3); in the main manuscript we show the results for windows containing 3,000 variable sites. We plotted the measures of π, *F_st_*, and *d_xy_*along the genome using the *manhattan* function from the qqman R package (v.0.1.4; Turner, 2014). The regions of increased or decreased divergence (or nucleotide diversity) were defined as the windows in the upper or lower 1% of the *F_st_*, *d_xy_*, and π distributions.

### Topology discordance in the nuclear genome

We reconstructed phylogenetic trees for non-overlapping sliding windows along the genome with PhyML (v.3.3.2; Guindon et al., 2010) using *phyml_sliding_windows.py* (https://github.com/simonhmartin/genomics_general) on the SNPs dataset including the *A. percula* individual, which was used to root the topologies. We used windows that were identical to those described above. We summarized the topologies by calculating the exact weightings of all possible subtrees using *twisst* (method “complete”, 10,000 sampling iterations; Martin & Van Belleghem, 2017). The individuals were grouped by species (species-level analysis) or populations (population-level analysis; see Supplementary Information S2). Plots were produced in R with *plot_twisst*.*R* provided in *twisst*.

### Tests for ancestral admixture

The cytonuclear incongruence and the topological inconsistencies along the genome showed that *A. sandaracinos* is occasionally phylogenetically closer to *A. perideraion* than to *A. akallopisos*. We investigated whether this disparity resulted from past hybridization events rather than from incomplete lineage sorting (ILS) by testing for signals of introgression between these two species using ABBA-BABA tests (Green et al., 2010). We computed the Patterson’s *D* statistic for a genome-wide estimation of admixture (Green et al., 2010; Durand et al., 2011). Briefly, we estimated allele frequencies at each site for each population with *freq.py* (https://github.com/simonhmartin/genomics_general). We considered *A. percula* as the outgroup and *A. perideraion* and *A. sandaracinos* as potentially hybridizing populations, and we calculated the *D* statistics with the equation (2) reported in Durand et al. (2011; Supplementary Information S3). We applied a block jackknife procedure to test the significance of the *D*statistic using *jackknife.R* (https://github.com/simonhmartin/genomics_general). To ensure independent blocks, we set a block size of 1Mb, resulting in 903 blocks. We verified that the signal of introgression was consistent in all chromosomes and independent of the geographic origin of the populations (Supplementary Information S3).

We estimated the proportion of admixture with the *f* statistics (Durand et al., 2011). We approximated the expected excess of ABBA over BABA sites under complete admixture by setting two *A. perideraion* populations (either geographic or random populations) as the hybridizing populations and estimating the site proportions as described above (Supplementary Information S3). We obtained the standard error and 95% confidence interval of *f* by applying a block jackknife approach as described above.

We identified candidate genomic regions of introgression (CRI) along the genome between *A. perideraion* and *A. sandaracinos* by estimating the *f_d_* statistics (Martin et al., 2015) with *ABBABABAwindows.py* (https://github.com/simonhmartin/genomics_general; Supplementary Information S3). We used windows of 3,000 variable sites but removed windows containing less than 100 biallelic SNPs to avoid stochastic errors in the *f_d_* estimation (Martin et al., 2015). We defined CRI as the windows in the top 5% of the genome-wide *f_d_* distribution. This threshold was set based on the estimated genome-wide admixture proportion between *A. sandaracinos* and *A. perideraion* (i.e., 5.5% of admixed genomes). We visually verified that the regions of high *f_d_* also supported the alternative mitochondrial topology. As introgressed regions generally show lower absolute genetic divergence (*d_xy_*; Smith & Kronforst, 2013; Martin et al., 2015), we tested for significant differences in *d_xy_* between the CRI and the rest of the genome for each combination of species (Supplementary Information S4). The presence of additional signals of admixture, besides that of *A. perideraion* and *A. sandaracinos*, was tested using TreeMix with 3- and 4-populations tests (v.1.13.; Pickrell & Pritchard, 2012; Supplementary Information S5).

### Gene content of the CRI and regions of increased/decreased divergence

We downloaded the structural gene annotation for 23,718 protein-coding genes of *A. percula* from the Ensembl database (release 99; https://www.ensembl.org) and expanded this functional annotation based on the available information from the *A. frenatus* reference genome (Marcionetti et al., 2018). This resulted in 17,179 annotated genes, of which 14,002 had *biological process* gene ontologies annotations (GOs; Supplementary Information S6). We performed GO enrichment analysis of the genes located within (or partially overlapping with) the CRI using the topGO package (parameters: Fisher’s exact tests, weight01 algorithms, minimum node size of 3; v.2.26.0; Alexa & Rahnenfuhrer, 2016) and contrasting them against all the annotated protein-coding genes of *A. percula*. Results were considered significant at *p-*values < 0.01. No corrections for multiple testing were performed, following recommendations from the topGO manual. An identical strategy was employed for the GO enrichment analysis of the genes located in the regions of increased/decreased genomic divergence between species.

### Tests of hybrid speciation, gene flow and demographic reconstruction

To infer the demographic scenario that best fitted the genomic data and test the hypotheses for the evolution of the skunk complex, we performed model comparison under the coalescent framework developed in fastsimcoal2 (v.2.6; Excoffier et al., 2013). We removed SNPs with missing data using vcftools (--max-missing 1; total SNPs kept: 16,547,283; v.0.1.15; Danecek et al., 2011). We treated the two populations of *A. akallopisos* from the WIO (Kenya, Mayotte) as a single population. We extracted the multidimensional folded site frequency spectra (SFS) for each population using easySFS (https://github.com/isaacovercast/easySFS), considering 10 samples per population (Supplementary Information S7). The multidimensional SFS were used to compare 16 distinct demographic models (Fig. 2A, Supplementary Information S7). We first compared a model of strict isolation with models of hybrid origin of *A. sandaracinos* followed by strict isolation or by asymmetric gene flow with *A. akallopisos* (Fig. 2A, a-c). We further compared these models with scenarios of ancestral asymmetric gene flow between *A. perideraion* and either the *A. akallopisos-A. sandaracinos* ancestor or *A. sandaracinos* (Fig. 2A, d-f). We then built models to investigate whether more recent gene flow (i.e., throughout the divergence of the species) on its own (Fig. 2A, g-i) or with ancestral gene flow (Fig. 2A, j-l), was likely. Finally, we explored the timing of the recent gene flow by comparing models with gene flow occurring only in the populations of the IAA (i.e., after the split of the allopatric populations; Fig. 2A n,p), and models where gene flow was only possible before the split of the allopatric populations (Fig. 2A, m, o). We did not include intraspecific gene flow in our demographic scenarios because it would have required the evaluation of additional and even more complex demographic models, which was out of the scope of this study. This choice could lead to an underestimation of divergence times if intraspecific gene flow is occurring between the populations (Leaché et al., 2014; Barley et al., 2018). It could also affect the estimation of population sizes, but we are only interested here in relative comparisons of *N_e_*, which have been shown to be valid despite the underestimation of gene flow (Leaché et al., 2014).

**Figure 2.**
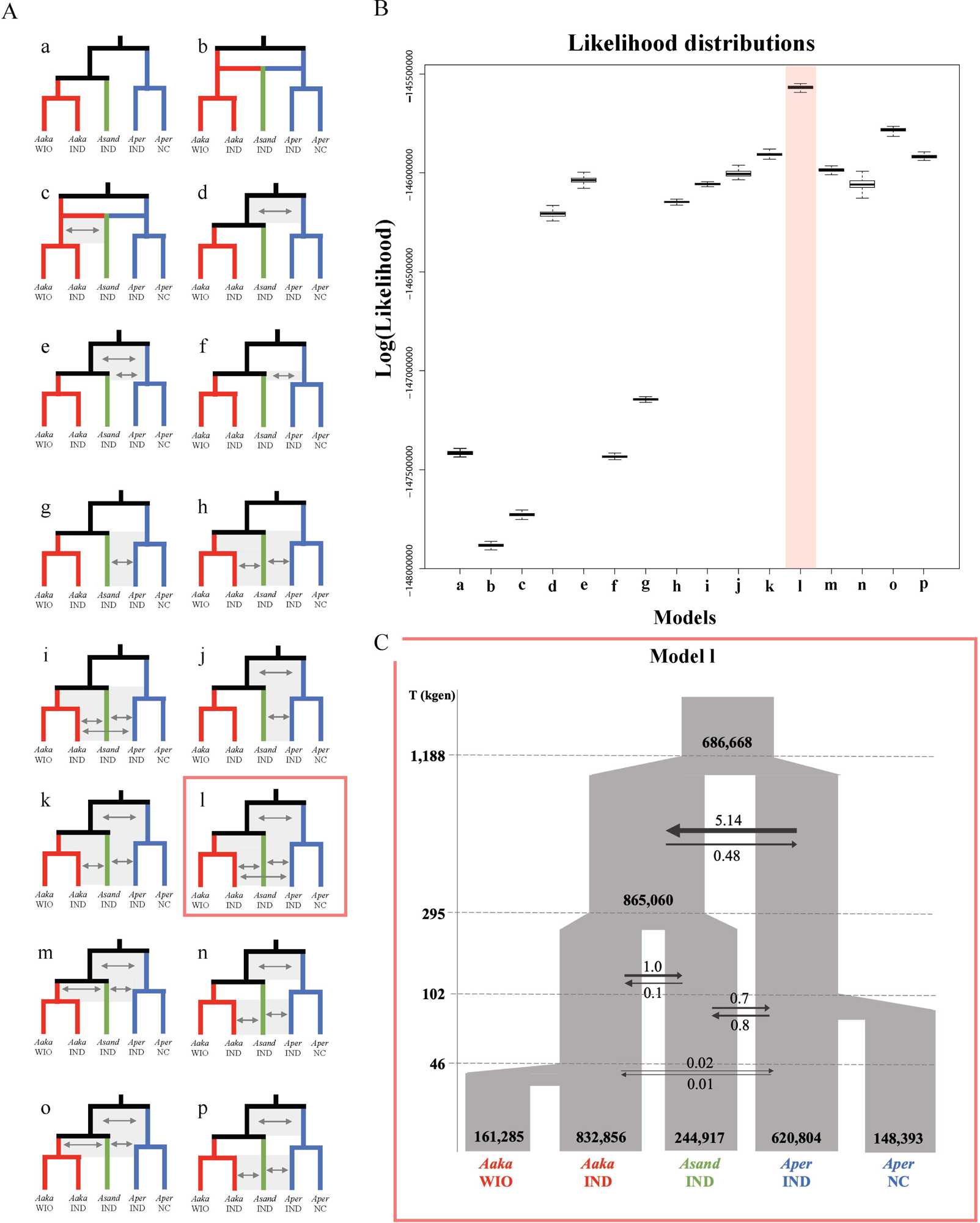
Schematic summaries of the 16 demographic models representing the possible diversification scenarios of the skunk complex (**A**) and the likelihood distributions obtained for them (**B**). Model l with the best-fit is highlighted in red in (**A)**, and a magnified version with the estimated parameters is reported in (**C**). The divergence times are reported in thousands of generations (kgen), and effective population sizes (*N_e_*, reported in black in the plot) are haploid sizes. Arrows represent gene flow between the populations, with numbers*10^-6^ corresponding to the estimated migration rates (*m*). Effective migration rates are calculated as *2N_e_m*. *Aaka, Asand* and *Aper* correspond to *A. akallopisos*, *A. sandaracinos* and *A. perideraion*, respectively. WIO, IND and NC correspond to the geographical origin of the populations (Western Indian Ocean, Indonesia, and New Caledonia, respectively). **A)** Arrows in the schematic models represent gene flow between the populations. Models of strict isolation (a), hybrid origin of *A. sandaracinos* (b, c) and divergence with different levels and timing of gene flow (d-p) were tested. **B**) Likelihood distributions were obtained based on 100 expected SFS, each approximated using 1 million coalescent simulations under the parameters that maximized the likelihood for each model. An overlap of distributions between models indicates no significant difference between the fit. Additional information on the analyses and results is provided in Supplementary Information S7.

We ran each model 50 times, performing 30 cycles of the expectation-conditional maximization (ECM) algorithm and considering 200,000 simulations to calculate the composite likelihood. We assumed a generation time of 5 years (as estimated for *A. percula*, Buston & Garcia, 2007) and set a mutation rate of 4.0×10^-8^ (derived from Delrieu-Trottin et al., 2017, see Supplementary Information S7). We identified the best-fitting demographic model based on both the rescaled AIC and the likelihood distributions obtained for each model (Supplementary Information S7).

For the best-supported model, we calculated confidence intervals of parameter estimates from 100 parametric bootstrap replicates by simulating SFS from the maximum composite likelihood estimates and re-estimating parameters each time (Excoffier et al., 2013; Lanier et al., 2015; Ortego & Sork, 2018). For each bootstrap replicate, we performed 30 independent runs with 200,000 simulations and 20 ECM cycles. We calculated point estimates of effective migration rates (i.e., “gene flow”) with 2*N_e_m* (with *N_e_* the haploid effective population sizes and *m* the migration to the given population), obtaining the number of gene copies exchanged each generation (Bourgeois et al., 2020). We defined limited (*N_e_m* = 0.01-0.1), weak (*N_e_m* = 0.1-1), and moderate (*N_e_m* = 1-10) gene flow as in Samuk & Noor (2022).

## RESULTS

### Whole Genome Sequencing, Mapping and SNP calling

The number of raw paired-end reads (PEs) for each individual ranged from ca. 29 million pairs to ca. 61 million pairs, with a median of ca. 48 million PEs per sample (Supplementary Table S4). After trimming low-quality regions, the number of PEs per sample ranged from 27 to 57 million, corresponding to an estimated raw coverage between 5.8X and 12.0X depending on the individual (Supplementary Table S4). Mapping to the *A. percula* reference genome resulted in the mapping of 84% to 88% of reads, with a final average coverage ranging between 4.6X and 10.2X (Supplementary Table S4).

The SNP calling strategy resulted in a total of 16,647,996 SNPs for the samples of the skunk complex, with an average SNPs density of 18.7 variants/kb. We observed regions of high and low SNPs density across all chromosomes (Supplementary Fig. S2). When the *A. percula* individual was included, the number of SNPs increased to 29,793,603 (Supplementary Table S5).

### Cytonuclear discordances in the skunk complex

We explored the phylogenetic relationship between the samples by reconstructing mitochondrial and nuclear phylogenetic trees. We found that both trees showed a clear separation of the three species, followed by a separation of populations depending on their geographical origin (Fig. 1B). Only the samples of *A. akallopisos* from the two populations of the WIO were not differentiated at both the nuclear and mitochondrial data.

Despite a clear separation of the species, the two genomes showed discordant topologies (Fig. 1B). At the nuclear level, *A. akallopisos* was sister to *A. sandaracinos*, while the latter was closer to *A. perideraion* at the mitochondrial level. This cytonuclear discordance was well supported, with bootstrap values higher than 0.95. We observed an additional inconsistency for one individual of *A. sandaracinos* (GB227), which grouped with its conspecifics at the nuclear level but clustered within the *A. perideraion* group from Indonesia in the mitochondrial tree (bootstrap support > 0.95; Fig. 1B).

### Overall clear divergence of the species and populations

We investigated the overall population structure by performing a PCA on the nuclear SNPs. We found that the first two axes split the three species into distinct clusters and explained 59.4% and 15.3% of the variance, respectively (Fig. 1C). The first axis separated the *A. sandaracinos* and *A. akallopisos* individuals from *A. perideraion*, while the second separated *A. akallopisos* from *A. sandaracinos*. The third and fourth components explained 3.8% and 2.3% of the variance, respectively. They separated the populations based on their geographical origin (Fig. 1C). The third axis split the Indonesian population of *A. perideraion* from the New Caledonian one, while the fourth axis divided the Indonesian population of *A. akallopisos* from the two populations in the Western Indian Ocean (WIO; Kenya and Mayotte). The admixture analysis also resulted in an overall separation of the individuals by species and geography. Indeed, the best number of ancestral populations was inferred to be K=3 and clustered the samples by species (Fig. 1D), but populations were separated with increasing K (K=4 and K=5; Supplementary Fig. S3).

In both the PCA and admixture results, individuals from the sympatric populations of the IAA never clustered together (Fig. 1C, 1D). Some signals of shared ancestry among species and populations were nevertheless observed in the admixture plots. For instance, with K=3 and K=4, samples of *A. akallopisos* from Indonesia showed a low proportion (6%) of shared ancestry with *A. sandaracinos*, while at K=2, a low proportion (13%) of *A. sandaracinos* ancestry was in common with *A. perideraion* (Fig. 1D, Supplementary Fig. S3).

Overall, measures of genomic divergence mirrored the PCA and admixture results. The averages *F_st_*calculated between all populations showed a clear species and population divergence, except for the two *A. akallopisos* populations of the WIO (Supplementary Information S1). Nevertheless, we identified high heterogeneity in the absolute (*d_xy_*) and relative (*F_st_*) genetic divergence across the genome (Fig 1E, Supplementary Fig. S4, S5). This result was stronger for the *F_st_* calculations but remained valid for *d_xy_*. The patterns were independent of the size of the sliding windows (Supplementary Table S3) or the considered populations within each species (Supplementary Fig. S4, S5). We looked for outlier regions of divergence (i.e., upper/lower 1% of the *F_st_*and *d_xy_* distributions) and found that all the windows of increased *d_xy_* and *F_st_* clustered in two regions of chromosome 18 for the comparisons *A. akallopisos* – *A. perideraion* and *A. sandaracinos – A. perideraion* (Fig 1E, Supplementary Fig. S5). These two regions also showed the lowest *d_xy_* between *A. akallopisos* and *A. sandaracinos*, and they were characterized by a reduced nucleotide diversity (π) in all populations (Fig 1E, Supplementary Fig. S4, S5, S6). In these two regions, extending from ca. 2.9 Mb to 3.5 Mb and from ca. 7.2 Mb to 16.9 Mb, we identified 408 functionally annotated genes, whose GO enrichment analysis resulted in 13 enriched GOs (*p-*value < 0.01, Supplementary Table S6). Among them, we observed terms associated with the regulation of behavior (GO:0050795), the development of endoderm (GO:0007492) and the morphogenesis of the epithelium (GO:1905332).

### Evidence from ancestral hybridization events and introgression in *A. sandaracinos*

We investigated the presence of topological inconsistencies along the nuclear genome of the skunk complex that may reflect ancestral gene flow before the split of populations within each species. Three different rooted topologies are possible linking the three species (Fig. 3A). The most frequent topology was the nuclear phylogenetic tree (“nuclear” topology in Fig. 3A), which had an average weighting higher than 80.0% and was fully supported in 74.8% of the windows (Fig. 3A, Supplementary Table S7). The topology of the mitochondrial phylogenetic tree (“mitochondrial” topology in Fig. 3A) was also represented in a relatively high proportion of the genome, being fully supported in 7.8% of the windows and having an average weighting of 13.0% (Fig. 3A, Supplementary Table S7). The last possible topology (“alternative” topology in Fig. 3A), characterized by *A. akallopisos* and *A. perideraion* being sister species, was fully supported in only 1.8% of the windows and had an average weighting of 7.0% (Fig. 3A, Supplementary Table S7). Overall, the three topologies were homogeneously distributed across the 24 chromosomes (Supplementary Fig. S7, Table S8). However, the weightings for the “mitochondrial” and “alternative” topologies were reduced on chromosome 18 (weightings of 4.6% and 1%, respectively) due to the presence of two large regions with full support for the “nuclear” topology (Fig. 3B, Supplementary Fig. S7, Table S8). These two regions coincided with two outlier regions of divergence (Fig. 1E).

**Figure 3.**
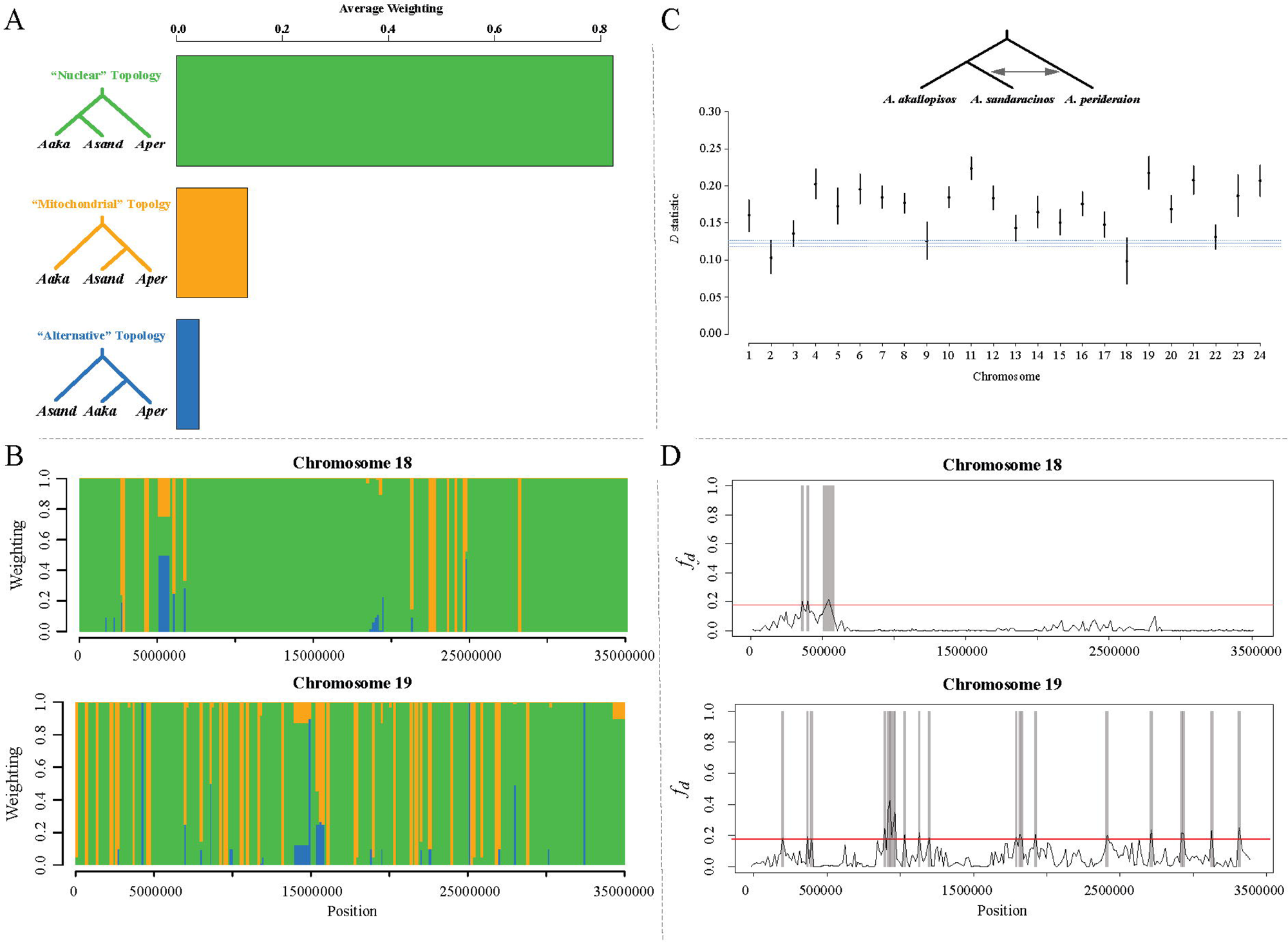
Whole-genome average weighting for each species-level topology (**A**) and examples of topological weighting across chromosomes 18 and 19 (**B**). **C**) The “mitochondrial” topology along the nuclear genome suggested ancestral gene flow between *A. sandaracinos* and *A. perideraion*, which was confirmed by the significantly positive values of Patterson’s *D* statistics. **D**) Examples of the *f_d_* distribution along chromosomes 18 and 19. Candidate regions of introgression (CRI) between *A. sandaracinos* and *A. perideraion* are reported in grey (**D**). (**A, B**) Results were obtained with *twisst*, for sliding windows of 3,000 variant sites. The weightings are determined by successively sampling a single member of each species and identifying the topology matched by the resulting subtree. Weightings are calculated as the frequency of occurrence of each topology in each window, and the average weightings are calculated genome-wide. Patterns observed on chromosome 19 are similar to those of the other chromosomes, while chromosome 18 shows two large regions of full support for the “nuclear” topology (green). (**C**) The *D* statistics ± s.e. are reported for each chromosome. Plotted in blue is the genome-wide estimate of *D* ± s.e..Standard errors were obtained with a block jackknife approach. All chromosomes show a *D-*statistic significantly deviating from zero, indicating introgression between *A. perideraion* and *A. sandaracinos*. (**D**) Red lines represent the 95th percentile of the genome-wide *f_d_* distribution, which was set as the cutoff for determining the CRI. Here also, patterns observed on chromosome 19 are similar to those of the other chromosomes, while chromosome 18 shows two regions of extremely low (< 0.01) *f_d_* and a reduced number of CRI.

The frequency of the “mitochondrial” topology along the nuclear genome (Fig. 3A), together with the cytonuclear discordance (Fig. 1B) and the admixture result for K=2 (Supplementary Fig. S3), suggested the presence of ancestral gene flow between *A. sandaracinos* and *A. perideraion*. This hypothesis was confirmed by significant and positive Patterson’s *D* statistics (0.171 ± 0.0041; Z-score of 41.9; *p*-value < 1E-045; Fig. 3C, Supplementary Table S9), with an overall proportion of admixture between the two species estimated to be 5.6% (*f* statistics=0.056, 95% CI [0.052-0.060]). The signal of admixture was detected in all 24 chromosomes, all of which showed significant *D* statistics (Fig. 3C, Supplementary Table S9) and the presence of candidate regions of introgression between the two species (CRI; upper 5% of the *f_d_* distribution; Fig. 3D, Supplementary Fig. S8). Chromosome 18 differed from the remaining chromosomes, showing the lowest values for the *D* statistics (0.1 ± 0.03, Z-score of 3.15; Fig. 3C, Supplementary Table S9) and containing two large regions with an extremely low *fd* (<0.01) and the absence of CRI (Fig. 3D).

We investigated the gene content of the CRI and identified 905 functionally annotated genes and 21 enriched GOs (*p*-value < 0.01, Supplementary Table S10). We observed terms associated with the sensory perception of a light stimulus (GO:0050953), the adult feeding behavior (GO:0008343), the neuropeptide signaling pathway (GO:0007218), as well as terms linked with the immune system (Supplementary Table S10).

### Weak evidence for ongoing extensive hybridization in the IAA

While the Indonesian populations of *A. perideraion*, *A. sandaracinos,* and *A. akallopisos* were differentiated from each other (Fig. 1C) and showed a relatively high genomic divergence (Supplementary Information S1 and Fig. S4), the admixture analysis suggested some level of shared ancestry between the Indonesian populations of *A. akallopisos* and *A. sandaracinos* (Supplementary Fig. S3). We investigated possible gene flow events in the IAA between the populations with TreeMix, which detected migration events between the Indonesian populations of *A. akallopisos* and *A. sandaracinos*, besides gene flow between *A. sandaracinos* and *A. perideraion* (see Supplementary Information S5).

We explored the presence of topological inconsistencies at the population level, with topologies grouping the three Indonesian populations together, which would suggest recent (i.e., after the population splits) gene flow in the IAA. We found an average topology weighting of only 3.4% for the topologies grouping the Indonesian populations of *A. akallopisos* and *A. sandaracinos*. We observed even lower average weighting when considering topologies with the Indonesian populations of *A. perideraion* and *A. sandaracinos* as sister species (1.8% average weighting), and with those of *A. perideraion* and *A. akallopisos* branching together (1.0%). In addition, we did not find any windows fully supporting any of these topologies. For more details, see Supplementary Information S2.

### Demographic reconstruction

We contrasted different diversification scenarios with or without gene flow (Fig. 2), potentially resulting in the observed patterns of admixture between the species. We obtained the highest likelihood for model l, in which gene flow happened between *A. perideraion* and the ancestor of *A. akallopisos + A. sandaracinos* and between all species throughout the diversification of the group (Fig. 2, Supplementary Information S7). Models of strict isolation and hybrid speciation resulted in the lowest likelihoods, and the additional models tested also showed reduced support compared to model l, with no overlap in the likelihood distributions (Fig. 2, Supplementary Information S7). Thus, we rejected the hypotheses of a hybrid origin of *A. sandaracinos*, of strict isolation during the group diversification, and the scenarios represented by the additional models.

Parameter estimates obtained for the best model reported the split between the three species to have occurred ca. 1.2 million generations ago and the one between *A. akallopisos* and *A. sandaracinos* ca. 0.3 million generations ago (Fig. 2, Supplementary Information S7). Considering a generation time of 5 years for clownfishes (Buston & Garcia, 2007), this corresponds to a divergence time of ca. 6 MYA and 1.5 MYA, respectively. The colonization of New Caledonia by *A. perideraion* was estimated to be ca. 509 kYA (101,809 generations), while the split of *A. akallopisos* populations appears to be more recent (ca 45,700 generations 228 kYA; Fig. 2C, Supplementary Information S7). The lowest population size estimates were observed for the New Caledonian population of *A. perideraion* and the *A. akallopisos* population from the WIO, while *A. sandaracinos* showed the smallest effective population size in the IAA (Fig. 2C, Supplementary Information S7). Estimates of the migration rates for the best model ranged between 1.3×10^-8^ and 5.14×10^-6^, depending on the timing and species considered (Fig. 2C, Supplementary Information S7). The highest rate (5.14×10^-6^) was observed from *A. perideraion* to the ancestor of *A. akallopisos + A. sandaracinos* and corresponded to an effective migration rate (2*N_e_m*) of 8.89 gene copies per generation. We obtained lower migration rates from *A. akallopisos* to *A. sandaracinos* (1×10^-6^) and from *A. perideraion* to *A. sandaracinos* (7.84×10^-7^), corresponding to effective migration rates (2*N_e_m*) of 0.49 and 0.38 gene copies per generation, respectively (Fig. 2C, Supplementary Information S7). The lowest migration rates were observed from *A. akallopisos* to *A. perideraion* (1.82×10^-8^) and vice versa (1.29×10^-8^) and corresponded to effective migration rates of 0.022 and 0.021 gene copies per generation, respectively (Fig. 2C, Supplementary Information S7).

## DISCUSSION

In addition to ecological speciation, gene flow between species and hybrid speciation were implicated with the divergence of the skunk group (Litsios et al., 2012; Litsios & Salamin, 2014). Here, we took advantage of the disjunct geographical distribution of the group to formally test these hypotheses and further investigate the mechanisms behind the diversification of the complex. First, we reject a hybrid origin of *A. sandaracinos*, but we show that moderate ancestral gene flow occurred from *A. perideraion* to the *A. akallopisos* + *A. sandaracinos* ancestor. Signals of introgression were detected in *A. sandaracinos* but absent in *A. akallopisos*. While we cannot exclude the role of natural selection in fixing the introgressed alleles in *A. sandaracinos*, the lower effective population size estimated for this species suggests a strong effect of genetic drift, which possibly allowed the introgressed DNA to reach fixation more extensively during the early stages of divergence. Second, we found evidence of weak gene flow between the three species throughout their diversification. Yet, the level of exchange was insufficient to impact the overall genetic divergence of the populations in the IAA, making it impossible to identify regions of increased differentiation potentially involved in ecological divergence. The absence of extensive gene flow among species despite their co-occurrence and their potential interactions within the same sea anemones hosts (Supplementary Table S1) suggests a role of host repartition and/or behavior-mediated reproductive barriers in maintaining the genetic identity of the species in sympatry. Finally, we identified two large regions of increased divergence on chromosome 18. This pattern indicates additional hybridization events with species outside the complex (Marcionetti & Salamin, 2023), with these regions being maintained by the disruption of recombination.

### Ancestral hybridization shaped the diversification of the skunk complex

Genomic comparisons of the skunk complex demonstrated that the three species were, overall, well-differentiated, with the separation of *A. perideraion* from *A. akallopisos* + *A. sandaracinos* estimated to have occurred around 6 MYA and the split of *A. akallopisos* and *A. sandaracinos* around 1.5 MYA. Nevertheless, we also confirmed cytonuclear discordance (Litsios & Salamin, 2014), with *A. perideraion* and *A. sandaracinos* being sister species at the mitochondrial level (Cowman et al., 2013; Frédérich et al., 2013; Rabosky et al., 2018). While the estimates for the divergence time of *A. perideraion* and the *A. akallopisos* + *A. sandaracinos* ancestor were consistent with previous studies (Rabosky et al., 2018), the lower divergence time for the split of *A. akallopisos* and *A. sandaracinos* obtained here compared to the literature (1.5 MYA vs 2.5 MYA from timetree.org; Kumar et al., 2017) suggests that the use of mitochondrial markers alone may have led to an overestimation of the divergence time between *A. akallopisos* and *A. sandaracinos* in previous studies.

Despite the overall clear genetic separation of the three species, topology discordances were also observed along the nuclear genome, and they suggested that the diversification of the skunk complex did not occur in strict isolation. Indeed, while such inconsistencies and cytonuclear discordance could also arise from ancestral polymorphism and incomplete lineage sorting (e.g., Toews & Brelsford, 2012; Lee-Yaw et al., 2019), we found here evidence of distinct hybridization events that occurred at different stages of the diversification of the skunk complex. The results of ABBA-BABA tests indicated past hybridization events between *A. perideraion* and *A. sandaracinos*, and demographic modeling allowed to identify a more complex scenario involving moderate gene flow (effective migration rate of 8.89 gene copies per generation) from *A. perideraion* to the *A. akallopisos* + *A. sandaracinos* ancestor and weak gene flow between the species throughout their evolution. These past hybridization events can account for the observed cytonuclear discordance. Indeed, hybridization likely led to the introgression and replacement of *A. sandaracinos* mitochondrial DNA by that of *A. perideraion* (i.e., mitochondrial capture, Toews & Brelsford, 2012; Perea et al., 2016), a phenomenon already documented in other Pomacentridae in the same geographic context (e.g., Bertrand et al., 2017). Similarly, following hybridization events, genomic regions of *A. perideraion* likely introgressed and got fixed by natural selection and/or genetic drift into *A. sandaracinos*, resulting in the (candidate) regions of introgression (CRI) present in *A. sandaracinos* genomes. The CRI contain genes with functions related to feeding behavior, perception of light, and immunity. These functions are broad and cannot be easily associated with shared ecological similarities in host usage and/or adaptive traits between *A. perideraion* and *A. sandaracinos*, which would suggest a role of natural or even sexual selection in the fixation of the regions (i.e., adaptive introgression; Hedrick, 2013).

While introgressed regions from *A. perideraion* were detected in *A. sandaracinos*, no clear signal of introgression was observed in *A. akallopisos*, as shown by the low support for the “alternative” topology (i.e., *A. akallopisos* sister to *A. perideraion*) along the nuclear genome (Fig. 3A). We observed therefore an imbalance of the introgression signal in *A. akallopisos* and *A. sandaracinos*, despite the stronger gene flow being detected between *A. perideraion* and the *A. akallopisos* + *A. sandaracinos* ancestor. This observation suggests that hybridization events between *A. perideraion* and the *A. akallopisos* + *A. sandaracinos* ancestor possibly occurred just before the speciation event of *A. akallopisos* and *A. sandaracinos*. Introgressed regions were then fixed at higher frequency in the newly-formed *A. sandaracinos* species, either through adaptive forces or through genetic drift, the latter being potentially stronger given the lower effective population size estimated for *A. sandaracinos* compared to *A. akallopisos* (Fig. 2C; Martin & Van Belleghem, 2017; Sagonas et al., 2019). Alternatively, the CRI of *A. sandaracinos* could have also originated from subsequent but weaker gene flow only concerning *A. sandaracinos* and *A. perideraion*. Nevertheless, such gene flow must have predated the divergence of *A. perideraion* populations, as the signal of introgression did not depend on the populations considered (but see individual GB227 discussed below for a noticeable exception).

### Population divergence and low migration rates in the IAA

When considering within-species genomic variation, we observed a clear separation of the populations by geography, especially for populations separated by large distances. Indeed, the divergence between the IAA populations of *A. perideraion* and *A. akallopisos* and the peripheral populations (i.e., NC and WIO, respectively) was relatively high, while within the WIO, the differentiation between *A. akallopisos* populations was low (Supplementary Figure S4). These results were consistent with restricted gene flow across large geographic distances, likely due to the low potential for long-range dispersal in clownfishes. In clownfishes, the larval stage lasts between 10 and 15 days (Fautin & Allen, 1997), which typically corresponds to a dispersal range of 5-100 km (Planes et al., 2009; Salinas-de-León et al., 2012). Dispersal as far as 400 km, however, has been documented for *A. omanensis* (Simpson et al., 2014). The connectivity between the IAA and New Caledonia (ca. 6,000 km) or the WIO (ca. 8,000 km) is therefore reduced, with intraspecific gene flow likely occurring only through a “stepping stones”-process, resulting in the population differentiation observed here. Within the WIO, the two populations of *A. akallopisos* (Kenya and Mayotte) were less differentiated due to the reduced geographical distance and the presence of the South Equatorial and the East African Coastal Currents (Figure 1 in Schouten et al., 2003), which can maintain increased connectivity between populations. We also observed that peripheral populations (WIO and NC) showed reduced *N_e_* and slightly reduced nucleotide diversity compared to their IAA equivalents (Fig. 2C, Supplementary Table S3). These patterns likely resulted from founder events and range expansion (Peter & Slatkin, 2013; Pierce et al., 2014; Braasch et al., 2019) and were in accordance with the central-marginal hypothesis (Eckert et al., 2008), further corroborating the IAA origin of the skunk complex (Litsios et al., 2014; Huyghe & Kochzius, 2017).

Demographic modeling estimated a weak interspecific gene flow between the IAA populations of the three species throughout their diversification (effective migration rates from 0.01 to 0.86, Fig. 2C; Samuk & Noor, 2022), with the lowest rates observed between the sympatric populations of *A. akallopisos* and *A. perideraion*. In comparison, effective migration rates higher than nine gene copies/generation were detected between highland and lowland bird forms of the Réunion grey white-eye, *Zosterops borbonicus*. Such gene flow resulted in the homogenization of the genome and the detection of loci involved in ecological adaptation (Bourgeois et al., 2020). Similarly, genome-wide homogenization of two *Ramphocelus* tanager species was obtained thanks to even higher (>20) migration rates, allowing the identification of loci involved in plumage color divergence (Luzuriaga-Aveiga et al., 2021). In the skunk complex, such high levels of gene flow are not occurring, and the populations of the three species remain clearly separated in the IAA (Fig. 1C,1D, Supplementary Information S2), preventing the genome homogenization that would be essential for uncovering peaks of high differentiation potentially linked with ecologically-important traits or with the speciation process.

Hybridization events involving members of the skunk complex are prevalent in the aquarium trade and were expected to be frequent also in nature (Steinke et al., 2009), likely facilitated by the possible cohabitation of different clownfish species within the same sea anemones hosts (Supplementary Table S1; Gainsford et al., 2015). Our findings revealed that gene flow between any species pair of the skunk complex is possible, but not extensive. Interspecific competition drives host repartition and specialization in regions of co-occurrence, pushing the different clownfish species to settle in distinct sea anemone host species (Garcia et al., 2023). Therefore, micro-habitat specialization might further reduce cohabitation and interactions among different species in the IAA. This could prevent hybridization events and maintain the clear genetic boundaries between species as observed in this study. The limited occurrence of hybrids in the wild could also be due to specific-specific behaviors that assist in maintaining reproductive isolation. In particular, the social structure consisting of a dominant breeding pair and subordinate non-breeding individuals is likely to reduce the opportunities for hybridization by maintaining a stable reproductive couple over longer periods (several years in some cases; Gainsford et al., 2015). There could also be strong mate selection with hybrid individuals facing difficulties in finding suitable mates, as they may not fit the preferences of individuals from either parent species. The weak gene flow observed in the skunk complex could, nevertheless, account for the small proportion of shared ancestry detected between the Indonesian populations of *A. akallopisos* and *A. sandaracinos* (Supplementary Fig. S3) and for the cytonuclear inconsistency observed for *A. sandaracinos* individual GB227, which was sister to the Indonesian *A. perideraion* individuals at the mitochondrial level (Fig. 1B). Similar to what we observed at the species level for *A. sandaracinos*, the retention of an allospecific mitochondrial genome in GB227 likely emerged from mitochondrial capture (Toews & Brelsford, 2012; Perea et al., 2016; Bertrand et al., 2017). Here, hybridization events between *A. perideraion* and *A. sandaracinos* must have occurred after the split of the New Caledonian population of *A. perideraion* and were then followed by subsequent backcrossing with *A. sandaracinos*. Selective backcrossing eventually erased the introgression signal from the nuclear DNA, resulting in the cytonuclear discordance observed for GB227 (Toews & Brelsford, 2012; Perea et al., 2016; Bertrand et al., 2017).

### Divergence on chromosome 18

Chromosome 18 showed peculiar patterns of differentiation in the skunk complex, with two large regions (from ca. 2.9 Mb to 3.5 Mb and from ca. 7.2 Mb to 16.9 Mb) showing reduced genetic divergence between *A. sandaracinos* and *A. akallopisos* but increased divergence with *A. perideraion* (Fig. 1E, Supplementary Fig. S4, S5). These results are consistent with recent findings of ancestral hybridization events across the clownfish group, which revealed past hybridization between the ancestor of *A. akallopisos* + *A. perideraion* and some ancestral species outside the skunk complex (Marcionetti & Salamin, 2023). In *A. akallopisos*, two analogous regions on chromosome 18 had maintained the signal of introgression, likely through genomic inversions that disrupted recombination (Marcionetti & Salamin, 2023). This hypothesis is further supported in this study by the reduced nucleotide diversity observed in these regions (Supplementary Fig. S6) and the absence of introgression from *A. perideraion* to *A. sandaracinos* (Fig. 3D). Genomic inversions are often responsible for the persistence of supergenes (e.g., Zinzow-Kramer et al., 2015; Küpper et al., 2016; Branco et al., 2018). In these regions, we found genes potentially involved in the ecological preference of the species, with, for instance, enrichment for genes associated with the regulation of behavior. Nevertheless, the role played by these regions - if any - in species divergence cannot be readily determined without further investigations.

## DATA AVAILABILITY AND BENEFIT-SHARING

Raw Illumina reads are deposited in the SRA (BioProject PRJNA1022585, accessible upon acceptance of the manuscript) and mitochondrial assemblies will be made available in the Zenodo repository upon acceptance of the manuscript.

Benefits Generated: Benefits from this research accrue from the sharing of our data and results on public databases as described above. All fieldwork was performed in agreement with local regulations and in collaboration with local entities (University of Mayotte; University of New Caledonia; Kenya Marine and Fisheries Research Institute; Universitas Hasanuddin, Indonesia) following the “Access and Benefit Sharing” (ABS) principles of the Nagoya Protocol established by the Convention of Biological Diversity (CBD). We thank the local authorities for permits to collect samples and help with field logistics in Indonesia, Mayotte, Kenya, and New Caledonia. Research permits number are the following: Mayotte: 06/UTM/2016; New Caledonia: N°60912-895-2017/JJC; Kenya: NCST/RRI/12/1/BS/250; Indonesia: 55/PSTK/UH/XI/04.

## ACKNOWLEDGEMENTS

We thank the local authorities of Indonesia, Kenya, Mayotte, and New Caledonia for the permits to collect samples and their help with field logistics. A special thanks goes to the Quesnel family for the help and logistical support they provided during fieldwork in New Caledonia. We also thank the staff at the Lizard Island Research Station for fieldwork support, and acknowledge the Dingaal, Ngurrumungu and Thanhil peoples as traditional owners of the lands and waters of the Lizard Island region. We are appreciative of the Lausanne Genomic Technology Facility for the sequencing and the DCSR computing infrastructure of the University of Lausanne for computational support/resources. Funding: University of Lausanne funds, Swiss National Science Foundation, Grant Number: 31003A-163428. FC was supported by an Australian Research Council (ARC) Discovery Early Career Research Award DE200100620 and Discovery Project DP180102363.

## AUTHOR CONTRIBUTIONS

JAMB and NS designed the study. AM and JAMB performed the research and analyses. FC, MK, LP, GFAD, SH, AM and JAMB contributed to the sample collection. AM wrote the manuscript. All authors read, made corrections, and approved the final version of the manuscript.

