## Supplementary Information and Figures for "Recurrent gene flow events shaped the diversification of the clownfish skunk complex"

- **Supplementary Information S1.** Average $F_{st}$ between populations and species.
- **Supplementary Information S2.** Population-level topological inconsistency analysis.
- **Supplementary Information S3.** Test for ancestral admixture. Complement on the methods used.
- **Supplementary Information S4:** Absolute genetic divergence ($d_{xy}$) for the candidate regions of introgression.
- **Supplementary Information S5.** Evidence for additional admixture events.
- **Supplementary Information S6.** Gene content of the candidate regions of introgression (CRI) and regions of increased/decreased divergence. Complement on the methods used.
- **Supplementary Information S7.** Test of hybrid speciation, gene flow and demographic reconstruction. Complement on methods and results.

Supplementary Tables

- **Table S1.** Interactions with sea anemone species for the clownfish species of the skunk complex. Data is obtained from Fautin & Allen (1997) and Litsios et al. (2012).
- **Table S2.** Information on the sampled individuals (sample name, population, species, sample location).
- **Table S3.** Average nucleotide diversity $\pi$, relative ($F_{st}$) and absolute ($d_{xy}$) genetic divergence for each population and pairwise comparison, for different window sizes.
- **Table S4.** Sequencing and mapping statistics. Statistics for raw reads, processed reads, mapping, and processed mapping on *A. percula* reference.
- **Table S5.** Total number of SNPs (A) and number of SNPs at each filtering step (B).
- **Table S6.** GO enrichment results for the two outlier regions of divergence observed on chromosome 18.
- **Table S7.** *Twisst* results for the species-level analysis for different window sizes.

- **Table S8.** *Twisst* results for the species-level analysis in different chromosomes. Average weighting and percent of windows with complete support are reported. Results are reported for windows of 3,000 sites, but results were consistent for different window sizes.
- **Table S9.** *D*-statistics obtained for the whole-genome and for each chromosome. Standard error (SE) and Z scored were obtained by a block jackknife approach.We considered different *A. perideraion* populations as P3 (i.e., hybridizing population) and results for each population are reported.
- **Table S10.** GO enrichment results for the candidate regions of introgression (CRI) between *A. sandaracinos* and *A. perideraion*.

Supplementary Figures (reported below in the document)

- **Figure S1.** Occurrence points of the three species of the skink complex considered in the study (*A. akallopisos, A. perideraion, A. sandaracinos*) obtained from GBIF.org.
- **Figure S2.** SNPs density across chromosomes.
- **Figure S3.** Admixture results for the individuals of the skunk complex obtained from PCAngsd.
- **Figure S4.** Absolute genetic divergence ($d_{xy}$) along the genome for all pairwise comparisons.
- **Figure S5.** Absolute genetic divergence ($F_{st}$) along the genome for all pairwise comparisons.
- **Figure S6.** Absolute nucleotide diversity (π) along the genome for all populations
- **Figure S7.** *Twisst* results along the chromosomes for the species-level analyses.
- **Figure S8.** $f_d$ distribution along the chromosomes for the test of introgression between *A. perideraion* and *A. sandaracinos* individuals.

**Supplementary Information S1. Average $F_{st}$ between populations and species.**

<u>Methods</u>

To investigate the overall genomic differentiation between the populations and species, we calculated the average $F_{st}$ for each pair of populations. This was done on the SNPs dataset for the skunk complex using vcftools (v.0.1.15; Danecek et al., 2011), which calculates $F_{st}$ as in Weir and Cockerham (1984). This estimation of $F_{st}$ was shown to be robust to low sample size when a large number of markers is available (Willing et al., 2012).

<u>Results</u>

The average $F_{st}$ between each pair of populations showed a low level of differentiation between populations of the same species. The $F_{st}$ between *A. akallopisos* populations ranged from 0.009 in the WIO to 0.074 and 0.075 between those from Indonesian and WIO (Figure 1). A similar level of differentiation was observed between the two populations of *A. perideraion* ($F_{st}$ of 0.11; (Figure 1). The divergence for populations of different species increased to $F_{st}$ values of ca. 0.3 for *A. sandaracinos* - *A. perideraion* and of ca. 0.5 for the comparison of *A. perideraion* with the two other species (Figure 1). Despite a slight decrease in $F_{st}$ when comparing populations of different species within the IAA compared to allopatric populations, the differentiation between the species remains high in the IAA (Figure 1).

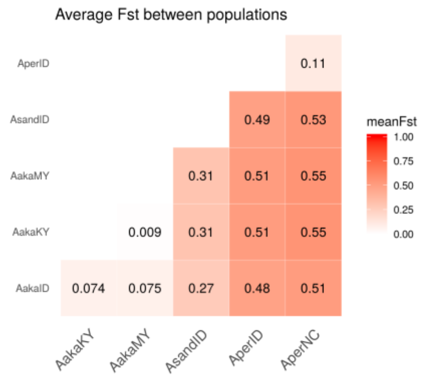


**Figure 1: Average $F_{st}$ obtained for all population pairs.** $f_{st}$ was calculated as in Weir and Cockerham (1984). AakaKY, AakaMY, AakaID correspond respectively to the *A. akallopisos* populations from Kenya, Mayotte and Indonesia. AperID and AperNC correspond respectively to the *A. perideraion* populations from Indonesia and New Caledonia. AsandID corresponds to the population of *A. sandaracinos* from Indonesia.

**Supplementary Information S2. Population-level topological inconsistency analysis.**

Methods

Population-level topological inconsistency analysis was performed similarly to the species level analysis (reported in the main text). After obtaining the phylogenetic trees for non-overlapping sliding windows with PhyML (v.3.3.2; Guindon et al., 2009), we performed *twisst* analyses (Martin & Van Belleghem, 2017, grouping individuals by populations. As the two populations of *A. akallopisos* from the Western Indian Ocean showed low divergence compared to the rest of the dataset (see also Supplementary Information S1), we merged them in the population-level analysis to decrease the number of possible topologies. Plots were produced in R with the script *plot_twisst.R* provided in *twisst*.

Results

We compared the topologies connecting the six populations along the genome to test for any evidence of introgression in sympatric populations within the IAA. We tested the 105 possible rooted binary trees existing for five populations. Out of the 105 trees, 96 of them showed values of average weighting measured lower than 1%, and we did not consider them further. Within the remaining nine topologies, only five were fully supported (weighting of 1) in at least two windows (Figure 1 and Table 1), and they together represented 78.1% of the windows across the genome (Figure 1). Results were overall consistent when considering different window sizes.

The most represented topology (average weighting of 56.4 %) was the expected species tree, with the two populations of *A. akallopisos* and *A. perideraion* branching first, and with *A. sandaracinos* as sister species to *A. akallopisos* (topology 1 in Figure 1). The second most frequent topology (average weighting of 13.9 %) corresponded to the mitochondrial topology, which displayed *A. sandaracinos* as sister species to *A. perideraion* (topology 2 in Figure 1). The three remaining topologies were less frequent along the genome (Figure). Topologies 3 and 5 described the alternative topology with *A. perideraion* as sister species to *A. akallopisos*, while topology 4 was similar to topology 1 (Figure 1). No support for topologies with populations of the IAA branching together was observed in any of the windows.

**Table 1: Windows with complete support for the different topologies for the population-level analysis.** The corresponding topologies can be seen in Figure 1 Data for windows of 3,000 sites is reported.

| | # Windows | % Windows | Mb of genome |
|---|---|---|---|
| **Topology 1** | 2498 | 44.92 | 398.33 |
| **Topology 2** | 492 | 8.85 | 78.92 |
| **Topology 3** | 6 | 0.11 | 0.91 |
| **Topology 4** | 6 | 0.11 | 0.81 |
| **Topology 5** | 4 | 0.07 | 0.39 |

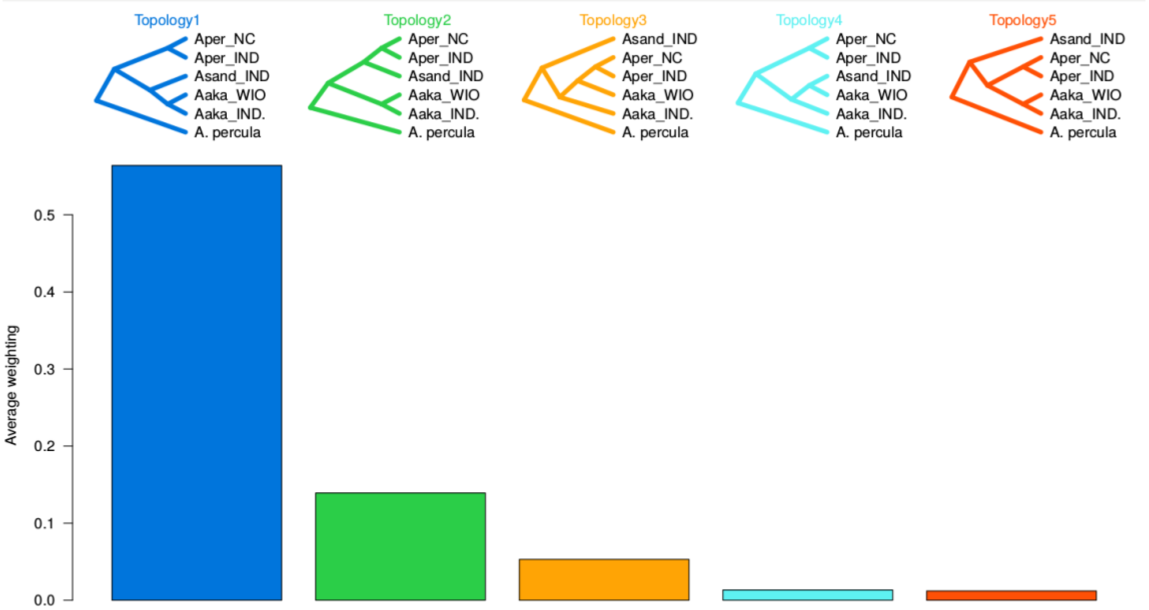


**Figure 1: Average proportion of subtrees matching each population-level topology (i.e., the weighting).** Topology 1 corresponds to the expected species topology, while topology 2 corresponds to mitochondrial topology. The names of the populations were abbreviated: Asand_IND for *A. sandaracinos* from Indonesia, Aper_IND and Aper_NC for *A. perideraion* from respectively Indonesia and New Caledonia, Aaka_IND and Aaka_WIO for *A. akallopisos* from respectively Indonesia and Western Indian Ocean (Mayotte and Keny). Results were obtained with *twisst*, for sliding windows of 3,000 variant sites.

When considering windows with complete support for each topology, we found that 44.9 % and 8.9 % of the windows showed full support for topology 1 and 2, respectively (Table 1). The three other topologies (topologies 3, 4, and 5) were fully supported in less than 1 % of the windows (Table 1). The remaining 45.9 % of the windows showed heterogeneous support, and similar results were obtained for different window sizes (Table 2). Such a proportion was considerably higher than what was observed in the species-level analysis (i.e., 15.8% of windows), likely due to the higher number of possible branching patterns (105 for the population level vs. 3 for the species level).

**Table 2**. ***Twisst* results of the population-level analysis for different window sizes.** The window size was based on the number of variants sizes in the skunk complex. A total of 105 possible rooted topologies were analyzed.

| | Window size (# of variants sites) | | | | | | | |
|---|---|---|---|---|---|---|---|---|
| | **1000** | **1500** | **2000** | **2500** | **3000** | **3500** | **4000** | **4500** |
| **# windows** | **16659** | **11110** | **8337** | **6671** | **5561** | **4769** | **4176** | 3712 |
| **Average Window size** | 53 kb | 80 kb | 107 kb | 133 kb | 160 kb | 186 kb | 213 kb | 240 kb |
| **# Topologies with no support** | 76 | 81 | 84 | 85 | 90 | 92 | 91 | 93 |
| **# Topologies > 1% overall weight** | 12 | 9 | 9 | 9 | 9 | 9 | 9 | 9 |
| **# Topologies with complete support** | 7 | 7 | 6 | 6 | 5 | 5 | 6 | 5 |
| **Overall % Topology 1** [a] | 47.5 | 50.9 | 53.1 | 54.5 | 56.4 | 57.2 | 58 | 58.9 |
| **Overall % Topology 2** [b] | 13.7 | 13.9 | 14.3 | 14.1 | 13.9 | 13.4 | 13.7 | 13.2 |
| **Overall % Topology 3** [c] | 5.3 | 5.6 | 5.3 | 5.3 | 5.3 | 5.6 | 5.5 | 5.6 |
| **Overall % Topology 4** [d] | 2.4 | 2 | 1.7 | 1.6 | 1.3 | 1.5 | 1.2 | 1.2 |
| **Overall % Topology 5** [e] | 1.3 | 1.2 | 1.2 | 1.2 | 1.2 | 1.1 | 1.1 | 1.2 |
| **Overall % weight of the 5 topologies** | 70.2 | 73.6 | 75.6 | 76.7 | 78.1 | 78.8 | 79.5 | 80.1 |
| **# Windows Complete support Topology 1** | 5702 | 4236 | 3432 | 2851 | 2498 | 2200 | 1965 | 1787 |
| **# Windows Complete support Topology 2** | 1236 | 891 | 706 | 585 | 492 | 409 | 384 | 327 |
| **# Windows Complete support Topology 3** | 24 | 22 | 17 | 10 | 6 | 5 | 4 | 1 |
| **# Windows Complete support Topology 4** | 74 | 37 | 23 | 18 | 6 | 6 | 3 | 4 |
| **# Windows Complete support Topology 5** | 15 | 9 | 7 | 6 | 4 | 4 | 4 | 3 |
| **% Windows Complete support Topology 1** | 34.23 | 38.13 | 41.17 | 42.74 | 44.92 | 46.13 | 47.05 | 48.14 |
| **% Windows Complete support Topology 2** | 7.42 | 8.02 | 8.47 | 8.77 | 8.85 | 8.58 | 9.20 | 8.81 |
| **% Windows Complete support Topology 3** | 0.14 | 0.20 | 0.20 | 0.15 | 0.11 | 0.10 | 0.10 | 0.03 |
| **% Windows Complete support Topology 4** | 0.44 | 0.33 | 0.28 | 0.27 | 0.11 | 0.13 | 0.07 | 0.11 |
| **% Windows Complete support Topology 5** | 0.09 | 0.08 | 0.08 | 0.09 | 0.07 | 0.08 | 0.10 | 0.08 |
| **% Windows Complete support** | 42.32 | 46.76 | 50.20 | 52.02 | 54.06 | 55.02 | 56.51 | 57.17 |
| **% Windows Mixed topologies** | 57.68 | 53.24 | 49.80 | 47.98 | 45.94 | 44.98 | 43.49 | 42.83 |
| **Complete support Topology 1 [Mb]** | 311.13 | 343.06 | 368.22 | 381.65 | 398.33 | 410.61 | 416.48 | 427.30 |
| **Complete support Topology 2[Mb]** | 65.50 | 72.20 | 76.39 | 78.38 | 78.92 | 78.11 | 83.52 | 79.54 |
| **Complete support Topology 3 [Mb]** | 1.21 | 1.63 | 1.72 | 1.25 | 0.91 | 0.86 | 0.73 | 0.24 |
| **Complete support Topology 4 [Mb]** | 3.52 | 2.56 | 2.08 | 1.94 | 0.81 | 0.76 | 0.53 | 0.72 |
| **Complete support Topology 5 [Mb]** | 0.55 | 0.45 | 0.45 | 0.55 | 0.39 | 0.49 | 0.53 | 0.44 |

a: (Apercul,(((Aaka_ID,Aaka_IO),Asand_ID),(Aper_ID,Aper_NC)))

b: (Apercul,((Aaka_ID,Aaka_IO),(Asand_ID,(Aper_ID,Aper_NC))))

c: (Apercul,((Aaka_ID,(Aaka_IO,(Aper_ID,Aper_NC))),Asand_ID))

e: (Apercul,(((Aaka_ID,Aaka_IO),(Aper_ID,Aper_NC)),Asand_ID))

f: (Apercul,((((Aaka ID,Aaka IO),Asand ID),Aper NC),Aper ID))

By considering all the possible topologies branching the Indonesian populations of *A. akallopisos* and *A. sandaracinos* as sister species, we obtained an average weighting of only 3.35%. We observed even lower average weighting when considering topologies with the Indonesian populations of *A. perideraion* and *A. sandaracinos* as sister species (1.8% average weighting) or with those of *A. perideraion* and *A. akallopisos* branching together (1.0%). Additionally, we did not find any windows fully supporting any of these topologies. The relationships among populations evidenced along the genome was mostly described by the topologies of the nuclear and mitochondrial phylogenies, with no evidence of topologies grouping populations from the same geographical location.

**Supplementary Information S3. Test for ancestral admixture. Complement on the methods used.**

We tested for signals of introgression between *A. perideraion* and *A. sandaracinos* by performing ABBA-BABA tests (Green et al., 2010). Given three populations and an outgroup with the relationship (((P1, P2), P3), O), ABBAs are the sites at which the derived allele B is shared between the non-sister taxa P2 and P3, whereas P1 carries the ancestral allele of the outgroup. Similarly, BABAs are the sites with the derived allele shared between P1 and P3, while P2 carries the ancestral allele. In the case of incomplete lineage sorting or recurrent mutation, the two types of sites should be equally abundant in the genome (Durand et al., 2011). In contrast, in the case of introgression between P2 and P3, a significant excess of ABBAs over BABAs should be observed. This excess can be tested using the Patterson's *D* statistic (Green et al., 2010; Durand et al., 2011) or the *fd* statistic (Martin et al., 2015). The *D*-statistic was designed to provide a genome-wide estimate of admixture, while *fd* provides a point estimate of the admixture proportion at specific loci (Martin et al., 2015).

***Genome-wide signal of admixture***

We first investigated genome-wide signals of introgression between species by estimating ABBAs and BABAs proportions. Counting ABBA and BABA sites in case of multiple samples per population can lead to the removal of a large amount of data, as the alleles that are not shared by all samples in each population cannot be considered. To avoid this problem, we can calculate the allele frequencies at each position for each population. Given the allele frequency of the derived allele (*pi*) and the ancestral allele (1-*pi*) in each population i (i=1, 2, 3 or O), ABBA and BABA proportions can then be estimated as follow:

$$ABBA = (1-p1)*p2*p3*(1-pO)$$

$$BABA = p1*(1-p2)*p3*(1-pO)$$

We obtained alleles frequencies for each population of the skunk complex and *A. percula* with the script *freq.py* (available from https://github.com/simonhmartin/genomics_general). We considered all the samples of the same species as a single population. We defined P1, P2, P3 as *A.*

*akallopisos*, *A. sandaracinos,* and *A. perideraion*, respectively, and *A. percula* was used as the outgroup. We calculated ABBAs and BABAs proportions based on the allele frequencies (as reported above) and estimated the *D*-statistic according to equation 1 in Durand et al. (2011). The equation is the following: $D=\frac{\sum ABBA-\sum BABA}{\sum ABBA+\sum BABA}$. By integrating the calculation of ABBA and BABA sites with allele frequencies (see above) in this equation, we obtained equation 2 in Durand et al. (2011), to which we refer in the main text.

To test for the significance of the *D*-statistic, we applied a block jackknife approach, which consisted of computing a "pseudo-mean" genome-wide *D*-statistics by iteratively excluding defined blocks of the genome. The standard error was then estimated by taking the difference between the mean genome-wide *D* and the "pseudo-mean" computed when the block is omitted. We computed the standard error and associated Z-score using the *jackknife.R* script (available from *https://*github.com/simonhmartin/genomics_general). To ensure independent blocks, we set a block size of 1Mb, resulting in 903 blocks.

We tested if the introgression signal was dependent on the geographical origin of the *A. perideraion* populations by setting P3 either as the *A. perideraion* population from Indonesia or the one from New Caledonia and repeated the analyses described above. We finally investigated whether all chromosomes showed evidence of introgression by applying the same procedure described above but estimating the *D*-statistic and associated standard error and Z-score for each chromosome independently.

***Estimation of the admixture proportion***

We estimated the proportion of admixture by estimating the *f* statistic (Durand et al., 2011). In this approach, we compared the observed excess of ABBA over BABA sites to the excess expected under complete admixture. To approximate the expectation under complete admixture, we set the populations of *A. perideraion* from New Caledonia and Indonesia as P2 and P3, respectively. ABBA and BABA proportions were then estimated in this setting (setting "B"), and we compared with the proportions observed with *A. sandaracinos* as P2 (setting "A"). The *f* statistic was then calculated as the ratio of the ABBA and BABA frequencies difference between settings A and B. The variance of the *f* statistic was estimated with a block jackknife approach, as described above, to obtain the standard error and 95% confidence interval of *f*. To ensure that the separation of the *A. perideraion* populations by geography in setting "B" was not biasing the analysis, we also estimated *f* by randomly distributing *A. perideraion* individuals in P2 and P3.

***Detection of potentially introgressed regions***

To identify regions of admixture, we applied the ABBA-BABA test in a sliding windows approach along the genome. We estimated the *fd* (Martin et al., 2015) and *fdm* statistics (Malinsky et al., 2015) using the script *ABBABABAwindows.py* (available from *https://*github.com/simonhmartin/genomics_general). Again, *A. akallopisos*, *A. sandaracinos,* and *A. perideraion* were set as P1, P2, and P3, respectively. The same windows obtained for the estimation of population genomic metrics were used. To avoid stochastic errors in the estimation of *fd* because of a small number of SNPs per window (Martin et al., 2015), we removed windows that contained less than 100 biallelic SNPs.

We defined candidate regions of introgression (CRI) as the regions falling into the top 5% of the genome-wide distribution of *fd*. We used this threshold because the estimate of the genome-wide proportion of admixture was found to be 5.5% between *A. sandaracinos* and *A. perideraion*. We visually verified that the locations with high *fd* fell into genomic regions supporting the alternative mitochondrial topology.

Because introgressed regions generally show lower absolute genetic divergence ($d_{xy}$; Smith & Kronforst, 2013; Martin et al., 2015), we compared $d_{xy}$ between *A. perideraion* and *A. sandaracinos* obtained for each window (as calculated in S*ection 2.8* in the main text) with the obtained *fd* values. We tested for a significant difference between the $d_{xy}$ observed in the CRI and the rest of the genome with the Welch two-sample t-test in R. The test was performed both with $d_{xy}$ values for the comparison *A. sandaracinos* - *A. perideraion* Indonesia, and *A. sandaracinos* - *A. perideraion* New Caledonia. We additionally tested for differences between the $d_{xy}$ values in CRI and the rest of the genome for the comparisons *A. akallopisos* – *A. perideraion* and *A. sandaracinos* – *A. akallopisos*. This was also performed with the Welch two-sample t-test.

**Supplementary Information S4. Absolute genetic divergence ($d_{xy}$) for the candidate regions of introgression**

Methods

Introgressed regions generally show lower absolute genetic divergence ($d_{xy}$) compared to non-introgressed regions (Smith & Kronforst, 2013; Martin et al., 2015). Therefore, we investigated whether the candidate regions of introgression (CRI) between *A. sandaracinos* and *A. perideraion* showed lower $d_{xy}$ compared to the rest of the genome. For this, we used Welch two-sample t-tests to test for significant differences in $d_{xy}$ (computed with *popgenWindows.py*, see main text) between the CRI and the rest of the genome for each combination of species. Regions on chromosome 18 that behave differently than the other chromosomes were not considered to avoid biases in the tests.

Results

For the comparison between *A. perideraion* (Indonesia) and *A. sandaracinos*, we found that $d_{xy}$ was significantly lower for the CRI (M=0.34, SD=0.045) compared to the rest of the genome (M=0.46, SD=0.061; t(332)=-40.6, $p < 0.001$; Figure 1). Consistent results were obtained between *A. perideraion* from New Caledonia and *A. sandaracinos* (CRI: M=0.34, SD=0.046; non-CRI: M=0.46, SD=0.061; t(330)=-40.3, $p < 0.001$).

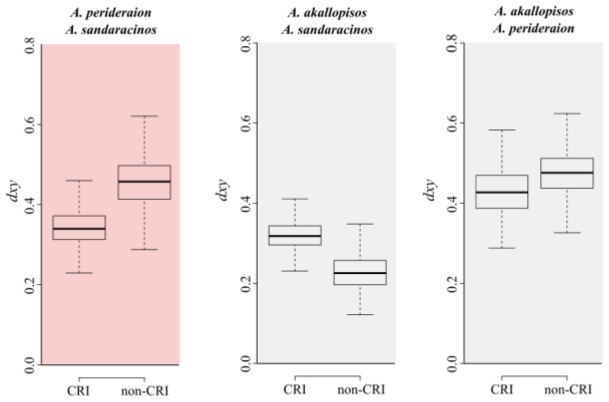


**Figure 1.** Comparisons of the absolute genetic divergence ($d_{xy}$) of the candidate regions of introgression (CRI, between *A. perideraion* and *A. sandaracinos*) and the rest of the genome (non-CRI) for each species pair. The CRI between *A. perideraion* and *A. sandaracinos* were defined as the regions in the top 5% of the $f_d$ distribution. The comparison of *A. perideraion* and *A. sandaracinos* is highlighted in red, as it depicts the $d_{xy}$ of the two introgressing species. Comparisons for Indonesian populations of the species are reported here, but comparable results were obtained for

each population.

In contrast, the CRI showed an increased $d_{xy}$ between *A. sandaracinos* and *A. akallopisos* compared to the rest of the genome (CRI: M=0.32, SD=0.041; non-CRI: M=0.23, SD=0.043; t(308)=36.4, $p < 0.001$; Figure 1), while they showed again a decreased $d_{xy}$ between *A. akallopisos* and *A. perideraion* (outliers regions: M=0.42, SD=0.060; non-outliers regions: M=0.47, SD=0.056; t(332)=-40.64, $p < 0.001$), despite this decrease was less pronounced (Figure 1).

The reduction of $d_{xy}$ between *A. sandaracinos* and *A. perideraion* in the CRI and its increase between *A. sandaracinos* and *A. akallopisos* suggest that gene flow occurred from *A. perideraion* to *A. sandaracinos*.

**Supplementary Information S5. Evidence for additional admixture events.**

Methods

In addition to the introgression signals between *A. sandaracinos* and *A. perideraion* detectable with the ABBA-BABA tests, other admixture events may have occurred during the diversification of the *akallopisos* group. Thus, to better understand and visualize the complexity of the ancestry of the skunk complex, we inferred patterns of population splits and mixtures with TreeMix (v.1.13.; Pickrell & Pickrell, 2012). This software computes a bifurcating maximum likelihood tree based on population allele frequencies, and it uses then a Gaussian approximation to estimate genetic drift between populations. Migration edges are then fitted between populations that are a poor fit to the tree model. The addition of migration events between branches is performed in stepwise iterations to maximize the likelihood until no further statistical significance is achieved.

For the TreeMix analysis, the SNPs dataset comprising *A. percula* was converted into TreeMix format using glactools (v.1.7.0; Renaud, 2018). We grouped samples according to populations or species. Because of the low divergence for *A. akallopisos* samples from the Western Indian Ocean (Mayotte and Kenya) compared to the rest of the dataset, we considered two populations as a single one. TreeMix was run for window sizes of 3,000 SNPs (-k parameter), using *A. percula* as outgroup (-root parameter) and by modeling 0 to 5 migration events (-m parameter). Different window sizes were tested and returned comparable results.

The *p*-values for the migration events computed by TreeMix are obtained by comparing a fixed graph structure with the migration event to the same graph without the migration. Consequently, significant *p*-values indicate that the hypothesized migration event significantly improves the fit of the data but do not necessarily mean that the tested migration is the correct one rather than another one between a different pair of populations. Thus, it is recommended to complement this analysis with less parameterized models (Pickrell & Pickrell, 2012). For this reason, we performed 3- and 4-populations tests (described in Reich et al., 2009). These tests rely on the *f3* and *f4* statistics, calculated based on differences in allele frequencies between populations. The *f3* method tests whether a target population is admixed from two source populations, while the *f4* method tests for the treeness of 4 species. In the case of admixture, these statistics are expected to be negative. The two tests were performed using scripts available in TreeMix (*threepop* and *fourpop* scripts, parameter -k set to 3,000). All possible combinations of three and four populations were tested. For the four-population tests, the analysis was also performed at the species level, i.e., by considering the four species as four populations.

Results

The TreeMix analyses identified two migrations events (Figure 1). The first was consistent with the ABBA-BABA test and corresponded to a migration from *A. perideraion* to *A. sandaracinos*. The second was a migration between *A. sandaracinos* and *A. akallopisos* from Indonesia. However, the weights for both migration events were low (0.05 for both migrations, Figure). The four-population test resulted in the detection of admixture between the different species (Table 1), in accord with the results obtained for the ABBA-BABA test. The three-population test resulted in the detection of mixture in *A. akallopisos* Indonesia when *A. akallopisos* from the Western Indian Ocean and either *A. sandaracinos* and or *A. perideraion* populations were considered as the source populations (Supplementary Table S15).

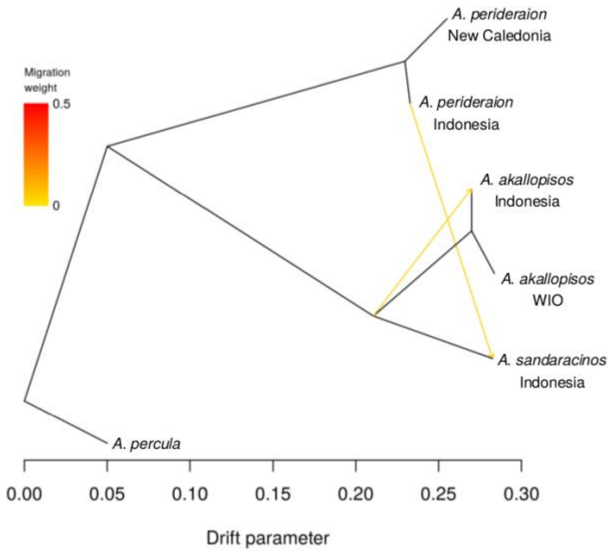


**Figure 1**: TreeMix Results. TreeMix maximum likelihood graph from whole-genome sequencing data, rooted with *A. percula*. Horizontal branch lengths are proportional to the amount of genetic drift that has occurred along the branch. Migration edges inferred using TreeMix are depicted as arrows colored by migration weight.

**Table 1**. **Results for the four-populations test**. The test was performed at the population level, testing all possible combinations of four populations. The test was also performed at the species level, i.e., considering all samples from the same species as a single population. Significant comparisons are reported in bold.

| Considered Trees | *f4* statistic | SE | Z score |
|---|---|---|---|
| **Population Level** | | | |
| AakaIO, AperID; Apercula, AsandID | -0.142587 | 0.00170924 | **-83.4216** |
| AakaIO, Apercula; AperID, AsandID | -0.152057 | 0.00156272 | **-97.3031** |
| AakaIO, AsandID; AperID, Apercula | -0.00947031 | 0.000278437 | **-34.0123** |
| AakaIO, AperNC; Apercula, AsandID | -0.143566 | 0.0017071 | **-84.0995** |
| AakaIO, Apercula; AperNC, AsandID | -0.153315 | 0.0015506 | **-98.8751** |
| AakaIO, AsandID; AperNC, Apercula | -0.00974935 | 0.000281783 | **-34.5988** |
| AakaID, AperID; Apercula, AsandID | -0.142129 | 0.00170206 | **-83.5041** |
| AakaID, Apercula; AperID, AsandID | -0.151178 | 0.00156904 | **-96.351** |
| AakaID, AsandID; AperID, Apercula | -0.00904933 | 0.000290377 | **-31.1641** |
| AakaID, AperNC; Apercula, AsandID | -0.143108 | 0.00170016 | **-84.173** |
| AakaID, Apercula; AperNC, AsandID | -0.15247 | 0.00155617 | **-97.9777** |
| AakaID, AsandID; AperNC, Apercula | -0.0093617 | 0.000290859 | **-32.1864** |
| AakaID, AakaIO; AperID, AperNC | 3.33E-005 | 1.53E-005 | 2.18181 |
| AakaID, AperID; AakaIO, AperNC | 0.396494 | 0.00253971 | 156.118 |
| AakaID, AperNC; AakaIO, AperID | 0.396461 | 0.00254108 | 156.02 |
| AakaID, AakaIO; AperID, Apercula | 0.000420985 | 9.09E-005 | 4.62984 |
| AakaID, AperID; AakaIO, Apercula | 0.216439 | 0.00129773 | 166.783 |
| AakaID, Apercula; AakaIO, AperID | 0.216018 | 0.00131442 | 164.345 |
| AakaID, AakaIO; AperID, AsandID | 0.00087917 | 0.000159047 | 5.52775 |
| AakaID, AperID; AakaIO, AsandID | 0.0743102 | 0.000725857 | 102.376 |
| AakaID, AsandID; AakaIO, AperID | 0.073431 | 0.000759544 | 96.6777 |
| AakaID, AakaIO; AperNC, Apercula | 0.000387647 | 8.32E-005 | 4.65778 |
| AakaID, AperNC; AakaIO, Apercula | 0.217697 | 0.00128816 | 168.998 |
| AakaID, Apercula; AakaIO, AperNC | 0.217309 | 0.00130324 | 166.745 |
| AakaID, AakaIO; AperNC, AsandID | 0.000845832 | 0.000151645 | 5.57771 |
| AakaID, AperNC; AakaIO, AsandID | 0.0745892 | 0.000731932 | 101.907 |
| AakaID, AsandID; AakaIO, AperNC | 0.0737434 | 0.000763705 | 96.5601 |
| AakaID, AakaIO; Apercula, AsandID | 0.000458185 | 9.36E-005 | 4.89733 |
| AakaID, Apercula; AakaIO, AsandID | 0.0648399 | 0.000510261 | 127.072 |
| AakaID, AsandID; AakaIO, Apercula | 0.0643817 | 0.000529061 | 121.691 |
| AakaID, AperID; AperNC, Apercula | -0.180055 | 0.0012755 | -141.165 |
| AakaID, AperNC; AperID, Apercula | -0.178764 | 0.00129173 | -138.391 |
| AakaID, Apercula; AperID, AperNC | 0.00129122 | 0.000151056 | 8.54791 |
| AakaID, AperID; AperNC, AsandID | -0.322184 | 0.00290445 | -110.928 |
| AakaID, AperNC; AperID, AsandID | -0.321872 | 0.00291494 | -110.421 |
| AakaID, AsandID; AperID, AperNC | 0.000312372 | 8.28E-005 | 3.77351 |
| AakaIO, AperID; AperNC, Apercula | -0.180443 | 0.00126626 | -142.501 |
| AakaIO, AperNC; AperID, Apercula | -0.179185 | 0.00128132 | -139.844 |
| AakaIO, Apercula; AperID, AperNC | 0.00125788 | 0.000152533 | 8.24657 |
| AakaIO, AperID; AperNC, AsandID | -0.32303 | 0.00290162 | -111.327 |
| AakaIO, AperNC; AperID, AsandID | -0.322751 | 0.0029111 | -110.869 |
| AakaIO, AsandID; AperID, AperNC | 0.000279035 | 8.26E-005 | 3.37777 |
| AperID, AperNC; Apercula, AsandID | -0.000978843 | 0.000130018 | -7.52849 |
| AperID, Apercula; AperNC, AsandID | 0.169715 | 0.00138903 | 122.182 |
| AperID, AsandID; AperNC, Apercula | 0.170693 | 0.00138457 | 123.283 |
| **Species Level** | | | |
| Aaka, Aper; Apercula, Asand | -0.0681325 | 0.000464436 | **-146.699** |
| Aaka, Apercula; Aper, Asand | -0.0732044 | 0.000424073 | **-172.622** |
| Aaka, Asand; Aper, Apercula | -0.0050719 | 9.71E-005 | **-52.2296** |

Aaka: All populations of *A. akallopisos*
AakaID: *A. akallopisos* from Indonesia
AakaWIO: *A. akkallopisos* from the Western Indian Ocean
Aper: All populations of *A. akallopisos*
AperID: *A. perideraion* from Indonesia
AperNC: *A. perideraion* from New Caledonia
Asand: All populations of *A. sandaracinos*
AsandID: *A. sandaracinos* from Indonesia

**Table 2. Results for the three-populations test**. The test was performed at the population level, testing all possible comparisons of three populations. The tested population corresponds to the population potentially arising from the admixture between the two source populations. Significant comparisons are reported in bold.

| Tested Population | Source Population 1 | Source Population 2 | *f3* statistic | SE | Z-score |
|---|---|---|---|---|---|
| *A. akallopisos* Indonesia | *A. akallopisos* WIO | *A. perideraion* Indonesia | **-0.00331915** | **0.000128358** | **-25.8586** |
| *A. akallopisos* Indonesia | *A. akallopisos* WIO | *A. perideraion* New Caledonia | **-0.00328581** | **0.000124721** | **-26.3453** |
| *A. akallopisos* Indonesia | *A. akallopisos* WIO | *A. sandaracinos* Indonesia | **-0.00243998** | **0.000106981** | **-22.8075** |
| *A. akallopisos* Indonesia | *A. perideraion* Indonesia | *A. perideraion* New Caledonia | 0.393175 | 0.00253231 | 155.264 |
| *A. akallopisos* Indonesia | *A. perideraion* Indonesia | *A. sandaracinos* Indonesia | 0.070991 | 0.000752661 | 94.32 |
| *A. akallopisos* Indonesia | *A. perideraion* New Caledonia | *A. sandaracinos* Indonesia | 0.0713034 | 0.000758085 | 94.0572 |
| *A. akallopisos* WIO | *A. akallopisos* Indonesia | *A. perideraion* Indonesia | 0.0171221 | 0.00024364 | 70.2762 |
| *A. akallopisos* WIO | *A. akallopisos* Indonesia | *A. perideraion* New Caledonia | 0.0170888 | 0.00023896 | 71.513 |
| *A. akallopisos* WIO | *A. akallopisos* Indonesia | *A. sandaracinos* Indonesia | 0.0162429 | 0.000176967 | 91.7852 |
| *A. akallopisos* WIO | *A. perideraion* Indonesia | *A. perideraion* New Caledonia | 0.413583 | 0.00248833 | 166.209 |
| *A. akallopisos* WIO | *A. perideraion* Indonesia | *A. sandaracinos* Indonesia | 0.0905531 | 0.000815066 | 111.099 |
| *A. akallopisos* WIO | *A. perideraion* New Caledonia | *A. sandaracinos* Indonesia | 0.0908321 | 0.000821575 | 110.559 |
| *A. perideraion* Indonesia | *A. akallopisos* Indonesia | *A. akallopisos* WIO | 0.399148 | 0.00251116 | 158.949 |
| *A. perideraion* Indonesia | *A. akallopisos* Indonesia | *A. perideraion* New Caledonia | 0.0026537 | 0.000278742 | 9.52028 |
| *A. perideraion* Indonesia | *A. akallopisos* Indonesia | *A. sandaracinos* Indonesia | 0.324838 | 0.0028767 | 112.92 |
| *A. perideraion* Indonesia | *A. akallopisos* WIO | *A. perideraion* New Caledonia | 0.00268704 | 0.000279887 | 9.60043 |
| *A. perideraion* Indonesia | *A. akallopisos* WIO | *A. sandaracinos* Indonesia | 0.325717 | 0.00287313 | 113.366 |
| *A. perideraion* Indonesia | *A. perideraion* New Caledonia | *A. sandaracinos* Indonesia | 0.00296607 | 0.00025606 | 11.5835 |
| *A. perideraion* New Caledonia | *A. akallopisos* Indonesia | *A. akallopisos* WIO | 0.422305 | 0.00240581 | 175.535 |
| *A. perideraion* New Caledonia | *A. akallopisos* Indonesia | *A. perideraion* Indonesia | 0.025844 | 0.000378303 | 68.3157 |
| *A. perideraion* New Caledonia | *A. akallopisos* Indonesia | *A. sandaracinos* Indonesia | 0.347716 | 0.00274291 | 126.769 |
| *A. perideraion* New Caledonia | *A. akallopisos* WIO | *A. perideraion* Indonesia | 0.0258107 | 0.000379076 | 68.0885 |
| *A. perideraion* New Caledonia | *A. akallopisos* WIO | *A. sandaracinos* Indonesia | 0.348562 | 0.00273887 | 127.265 |
| *A. perideraion* New Caledonia | *A. perideraion* Indonesia | *A. sandaracinos* Indonesia | 0.0255317 | 0.000357201 | 71.477 |
| *A. sandaracinos* Indonesia | *A. akallopisos* Indonesia | *A. akallopisos* WIO | 0.129327 | 0.000940497 | 137.509 |
| *A. sandaracinos* Indonesia | *A. akallopisos* Indonesia | *A. perideraion* Indonesia | 0.0558961 | 0.000448991 | 124.493 |
| *A. sandaracinos* Indonesia | *A. akallopisos* Indonesia | *A. perideraion* New Caledonia | 0.0555837 | 0.000441422 | 125.92 |
| *A. sandaracinos* Indonesia | *A. akallopisos* WIO | *A. perideraion* Indonesia | 0.0550169 | 0.000420177 | 130.937 |
| *A. sandaracinos* Indonesia | *A. akallopisos* WIO | *A. perideraion* New Caledonia | 0.0547379 | 0.000414881 | 131.937 |
| *A. sandaracinos* Indonesia | *A. perideraion* Indonesia | *A. perideraion* New Caledonia | 0.377768 | 0.00282853 | 133.556 |

**Supplementary Information S6. Gene content of the candidate regions of introgression (CRI) and regions of increased/decreased divergence. Complement on the methods used.**

We updated the functional annotation of the *A. percula* reference genome. Structural gene annotation for 23,718 protein-coding genes of *A. percula* was retrieved from the Ensembl database (release 99). Functional annotation was available for 15,875 protein-coding genes, with 9,453 of them linked to "biological process" gene ontologies (GO). To expand this information, we further annotated the whole *A. percula* proteome based on the protein-coding genes annotation of *A. frenatus* (Marcionetti et al., 2018) using reciprocal blast searches of the proteomes of *A. frenatus* and *A. percula* (blastp v.2.9.0; https://blast.ncbi.nlm.nih.gov/Blast.cgi). After filtering out poorly-supported hits (E-value cut-off of 10-6 and a minimum identity of 80%), we extracted best-reciprocal blast hits and considered them orthologous genes. We then annotated *A. percula* genes based on the functional annotation of the *A. frenatus* orthologs. We verified that the updated annotation was coherent with the original *A. percula* annotation. In the case of close-by genes with identical annotations, we only kept one gene. This allowed reducing the bias in the GO enrichment analysis arising from a potentially fragmented structural annotation. The extended annotation resulted in a total of 17,179 annotated genes, with 14,002 of them annotated with "biological process" GO.

We retrieved the genomic coordinates of the candidate introgressed regions (CRI; see *Section 2.10* in the main text) and the regions of increased genomic divergence between species (see *Section 2.8* in the main text), and we retrieved the genes located within these regions. Genes only partially overlapping the regions were also considered as candidate loci. For both analyses, we performed GO enrichment by contrasting the annotation of the genes in the regions of interest (i.e., CRI or higher divergence regions) against all the annotated protein-coding genes of *A. percula*. We used the TopGO package (v.2.26.0; Alexa & Rahnenfuhrer, 2016) available in Bioconductor (http://www.bioconductor.org), setting a minimum node size of 3. Fisher's exact tests with the weight01 algorithms were applied to examine the significance of enrichment in each analysis, with *p-values* < 0.01 considered significant. We present *raw p-values* instead of *p-values* corrected for multiple testing, following recommendations from the topGO manual.

**Supplementary Information S7. Test of hybrid speciation, gene flow and demographic reconstruction. Complement on methods and results.**

Methods

We retrieved the SNPs dataset without the *A. percula* sample and further removed positions with missing data using vcftools (--max-missing 1; v.0.1.15; Danecek et al., 2011). This resulted in a total of 16,547,283 SNPs (100,713 removed positions). We considered the two populations of *A. akallopisos* from the Western Indian Ocean as a single population. We extracted the multidimensional folded site frequency spectra (SFS) for each population using easySFS (https://github.com/isaacovercast/easySFS). We selected the projection that maximized the number of segregating sites per population using the –preview option of easySFS. This corresponded to 10 samples per population.

The multidimensional SFS were used to compare a total of 16 distinct demographic models (Supplementary Figure S2). We first compared a model of strict isolation (Figure 1A) with models where *A. sandaracinos* originated from the hybridization of *A. akallopisos* and *A. perideraion* (i.e., hybrid speciation) followed by strict isolation (Figure 1B) or by asymmetric gene flow between *A. akallopisos* and *A. sandaracinos* (Figure 1C). We further compared these models with scenarios of ancestral asymmetric gene flow between the *A. akallopisos-A. sandaracinos* ancestor and *A. perideraion* and/or between *A. sandaracinos* and *A. perideraion* (Figure 1D-F). We then built models to investigate whether more recent gene flow between the three species was likely, either alone (Figure 1G, 1H, 1I) or with ancestral gene flow (Figure 1J, 1K, 1L). In these scenarios, we modeled gene flow throughout the divergence of different pairs of Indonesian populations (*A. sandaracinos - A. perideraion,* Figure 1G, 1J; *A. sandaracinos - A. perideraion* and *A. akallopisos - A. sandaracinos*, Figure 1H, 1L; *A. sandaracinos - A. perideraion, A. akallopisos - A. sandaracinos* and *A. akallopisos - A. perideraion*, Figure 1I, 1L). Finally, we explored the timing of the recent gene flow by comparing models with gene flow only in populations from the IAA region (i.e., after the split of the allopatric populations of *A. akallopisos* and *A. perideraion* (Figure 1N, 1P), and models where gene flow was only possible before the split of the allopatric populations (Figure 1M, 1O).

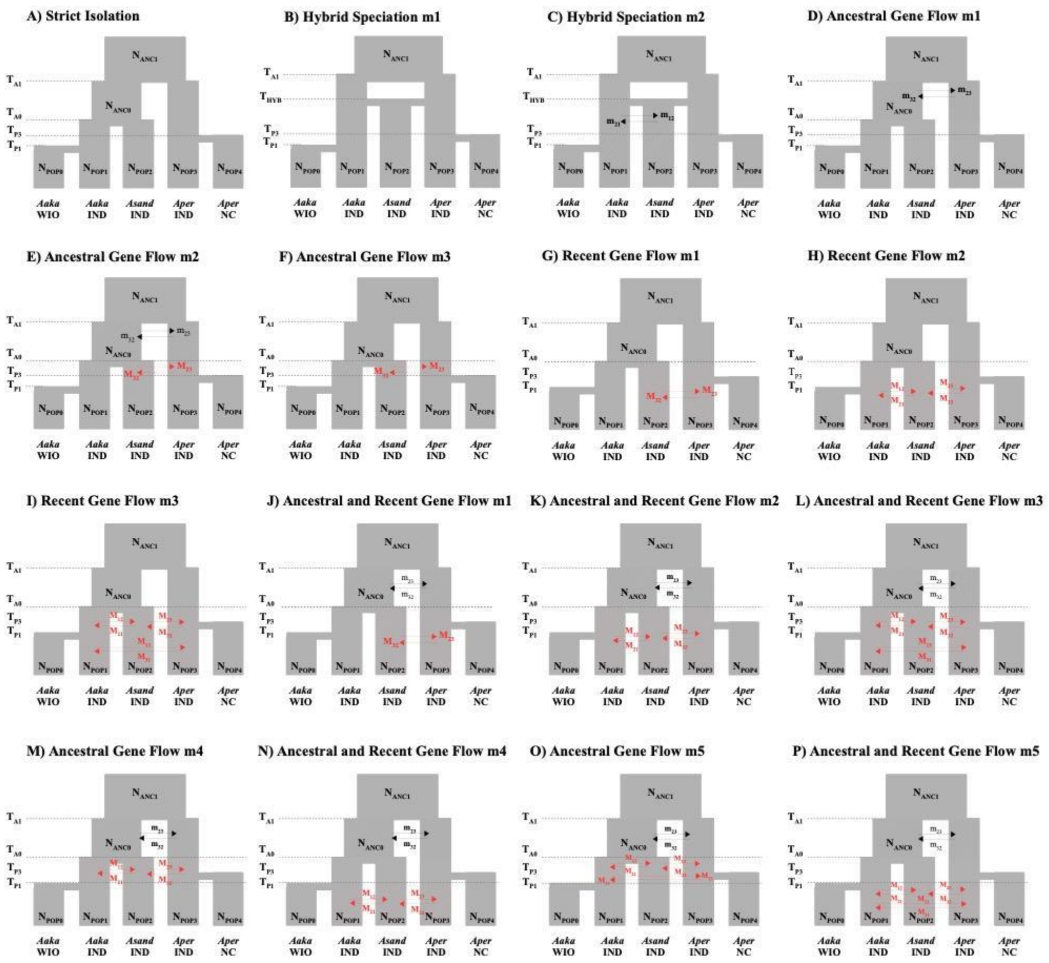


**Figure 1:** Summary of the 16 demographic models and their parameters evaluated using whole-genome SNPs. *Aaka*, *Asand*, and *Aper* correspond respectively to *A. akallopisos*, *A. sandaracinos*, and *A. perideraion*. WIO, IND, and NC correspond to the geographical origin of the population (respectively Western Indian Ocean, Indonesia, and New Caledonia). Gene flow between *A. perideraion* and the ancestor of *A. akallopisos* and *A. sandaracinos* is represented by black arrows. Red arrows correspond to more recent migrations (i.e., after the split between *A. akallopisos* and *A. sandaracinos*). We define "recent gene flow" when it is modeled to also occur in the present time. The exchanging populations were highlighted using hatching.

In all models, effective population sizes were estimated for each population and could vary at each splitting time. Parameters were estimated from the multidimensional SFS using fastsimcoal2 (v.2.6; Excoffier et al., 2013). We ran each model 50 times, performing 30 cycles of the expectation-conditional maximization (ECM) algorithm and considering 200,000 simulations to calculate the composite likelihood. We assumed a generation time of 5 years (as estimated for *A. percula*, Buston & Garcia, 2007) and set a mutation rate of 4.0x10-8 (obtained from the average expected mutation rate per site per year for nuclear genomes of reef fishes and assuming a generation time of clownfishes of 5 years; Delrieu-Trottin et al., 2017). For each model, we retained the set of parameters with the highest final likelihood as the best point estimate. We compared the expected and observed SFS with the SFStools.R (available from https://github.com/marqueda/SFS-scripts).

We identified the best-fitting demographic model based on the rescaled Akaike's information criterion (AIC; Akaike, 1974). AIC values should, however, be interpreted with caution as linked sites are present in our data (Excoffier et al., 2013). We thus also examined the likelihood distributions obtained based on 100 expected SFS, each approximated using 1 million coalescent simulations under the parameters that maximized the likelihood for each model. An overlap of these distributions between models indicates no significant difference between the fit of alternative models.

For the best-supported model, we calculated confidence intervals of parameter estimates from 100 parametric bootstrap replicates by simulating SFS from the maximum composite likelihood estimates and re-estimating parameters each time (Excoffier et al., 2013; Lanier et al., 2015; Ortego & Sork, 2018). For each bootstrap replicate, we performed 30 independent runs with 200,000 simulations and 20 ECM cycles. We used the parameter point estimates from the run with the highest likelihood of each bootstrapping replicate to compute the 95 percentile confidence intervals. We calculated points estimates of effective migration rates with 2*N*m (with N the haploid population sizes and m the migration to the given population), obtaining the number of gene copies exchanged each generation (as in Bourgeois et al., 2020). We defined limited ($N_em$ = 0.01-0.1), weak ($N_em$ = 0.1-1), and moderate ($N_em$ = 1-10) gene flow as in Samuk & Noor (2022).

Results

The best result obtained was the model "ancestral gene flow and recent gene flow m3" (Figure 1L, Figure 2). This model had the highest likelihood of all tested models (Figure 2A, Table 1). In this model, gene flow happened between *A. perideraion* and the ancestor of *A. akallopisos* – *A. sandaracinos,* and also between all species throughout the diversification of the group. The estimate of the parameters and the confidence intervals are reported in Table 2.

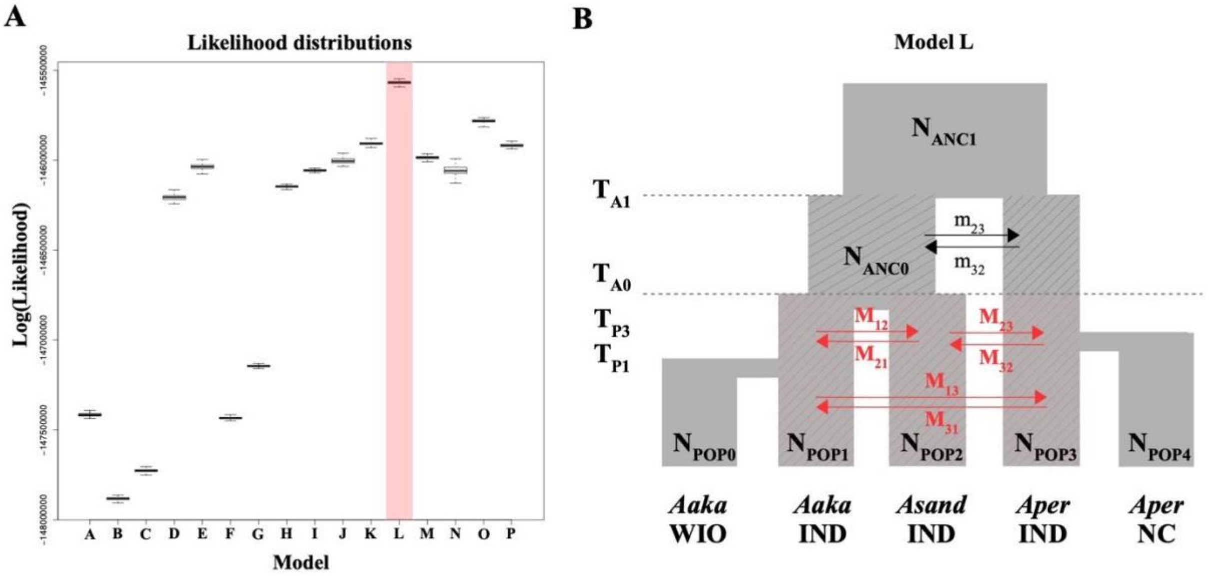


**Figure 2:** Likelihood distributions obtained for the 16 demographic scenarios (A) and the schematic representation of the resulting best model (B). A) The best resulting model (model L) is highlighted in red. All tested scenarios (A to P) are depicted in Figure 1. Likelihood distributions were obtained based on 100 expected SFS, each approximated using 1 million coalescent simulations under the parameters that maximized the likelihood for each model. An overlap of distributions between models indicates no significant difference between the fits. B) Gene flow between *A. perideraion* and the ancestor of *A. akallopisos* and *A. sandaracinos* is represented by black arrows. Red arrows correspond to more recent migrations (i.e., after the split between *A. akallopisos* and *A. sandaracinos*). We define "recent gene flow" when it is modeled to also occur in the present time. The exchanging populations were highlighted using hatching. Parameter estimates for the best model are reported in Supplementary Table S9.

**Table 1:** Likelihoods of the 16 demographic models compared with fastsimcoal2, and their AIC scores. Models are depicted in Figure 1.

| Model | Model Name | Log(likelihood) | # of Parameters | AIC | ΔAIC |
|---|---|---|---|---|---|
| L | Ancestral and Recent Gene Flow m3 | -145512945.9 | 19 | 291025930 | 0 |
| O | Ancestral Gene Flow m5 | *-145751170.2* | *19* | *291502378* | 476448 |
| P | Ancestral and Recent Gene Flow m5 | *-145858302.9* | *19* | *291716644* | 690714 |
| K | Ancestral and Recent Gene Flow m2 | *-145871281* | *17* | *291742596* | 716666 |
| J | Ancestral and Recent Gene Flow m1 | *-145935044.9* | *16* | *291870122* | 844192 |
| M | Ancestral Gene Flow m4 | *-145969250.1* | *17* | *291938534* | 912604 |
| N | Ancestral and Recent Gene Flow m4 | *-145974810.5* | *17* | *291949655* | 923725 |
| E | Ancestral Gene Flow m2 | *-145976769.4* | *15* | *291953569* | 927639 |
| I | Recent Gene Flow m3 | *-146019854.9* | *17* | *292039744* | 1013814 |
| H | Recent Gene Flow m2 | *-146113743.8* | *15* | *292227518* | 1201588 |
| D | Ancestral Gene Flow m1 | *-146116110.8* | *13* | *292232248* | 1206318 |
| G | Recent Gene Flow m1 | *-147119984.1* | *14* | *294239994* | 3214064 |
| A | Strict Isolation | *-147339577.1* | *11* | *294679176* | 3653246 |
| F | Ancestral Gene Flow m3 | *-147400513.9* | *13* | *294801054* | 3775124 |
| C | Hybrid Speciation m2 | *-147657871.7* | *13* | *295315769* | 4289839 |
| B | Hybrid Speciation m1 | -147827976.3 | 10 | 295655973 | 4630043 |

**Table 2:** Parameter estimates for the best model with ancestral and recent gene flow between species (model L). Population sizes are haploid sizes. Divergence times are reported in generations. The model is depicted in Figure 1L and Figure 2B. 95% Confidence Intervals (CI) were obtained by parametric bootstrap.

| Parameter | Estimate | 95% CI | Parameter | Estimate | 95% CI |
|---|---|---|---|---|---|
| $N_{POP0}$ | 161,285 | [161,285; 161,285] | $M_{13}$ | $1.82x10^{-8}$ | [$1.79x10^{-8}$; $1.86x10^{-8}$ |
| $N_{POP1}$ | 832,856 | [832,817.3; 832,894.6] | $M_{31}$ | $1.29x10^{-8}$ | [$1.25x10^{-8}$; $1.33x10^{-8}$ |
| $N_{POP2}$ | 244,917 | [244,917; 244,917] | $M_{21}$ | $8.59x10^{-8}$ | [$8.56x10^{-8}$; $8.63x10^{-8}$ |
| $N_{POP3}$ | 620,804 | [620,804; 620,804] | $M_{12}$ | $9.99x10^{-7}$ | [$9.99x10^{-7}$; $9.99x10^{-7}$ |
| $N_{POP4}$ | 148,393 | [148,393; 148,393] | $M_{23}$ | $7.08x10^{-7}$ | [$7.05x10^{-8}$; $7.11x10^{-8}$ |
| $N_{ANC0}$ | 865,060 | [865,060; 865,060] | $M_{32}$ | $7.84x10^{-7}$ | [$7.84x10^{-7}$; $7.84x10^{-7}$ |
| $N_{ANC1}$ | 686,668.3 | [686,636.9; 686,699.7] | $m_{32}$ | $5.14x10^{-6}$ | [$5.13x10^{-6}$; 5.143x10[illegible] |
| $T_{P1}$ | 45,697.27 | [45,628.75; 45,765.79] | $m_{23}$ | $4.82x10^{-7}$ | [$4.79x10^{-7}$; $4.84x10^{-7}$ |
| $T_{P3}$ | 101,809.9 | [101,750.6; 101,869.2] | | | |
| $T_{A0}$ | 294,654.8 | [294,519.2; 294,790.4] | | | |
| $T_{A1}$ | 1,188,136 | [1,184,580; 1,191,692] | | | |

**Supplementary Figure S1:** Occurrence points of the three species of the skink complex considered in the study (*A. akallopisos*, *A. perideraion*, *A. sandaracinos*) obtained from GBIF.org (GBIF Occurrence Download https://doi.org/10.15468/dl.rzpkl4). Occurrence data was downloaded on January, the 25th 2016, and filtered to keep only occurrence points with precise latitude and longitude information and in the expected range of the *Amphiprion* distribution. A total of 141, 223 and 31 occurrence points were obtained for *A. akallopisos*, *A. perideraion* and *A. sandaracinos*, respectively.

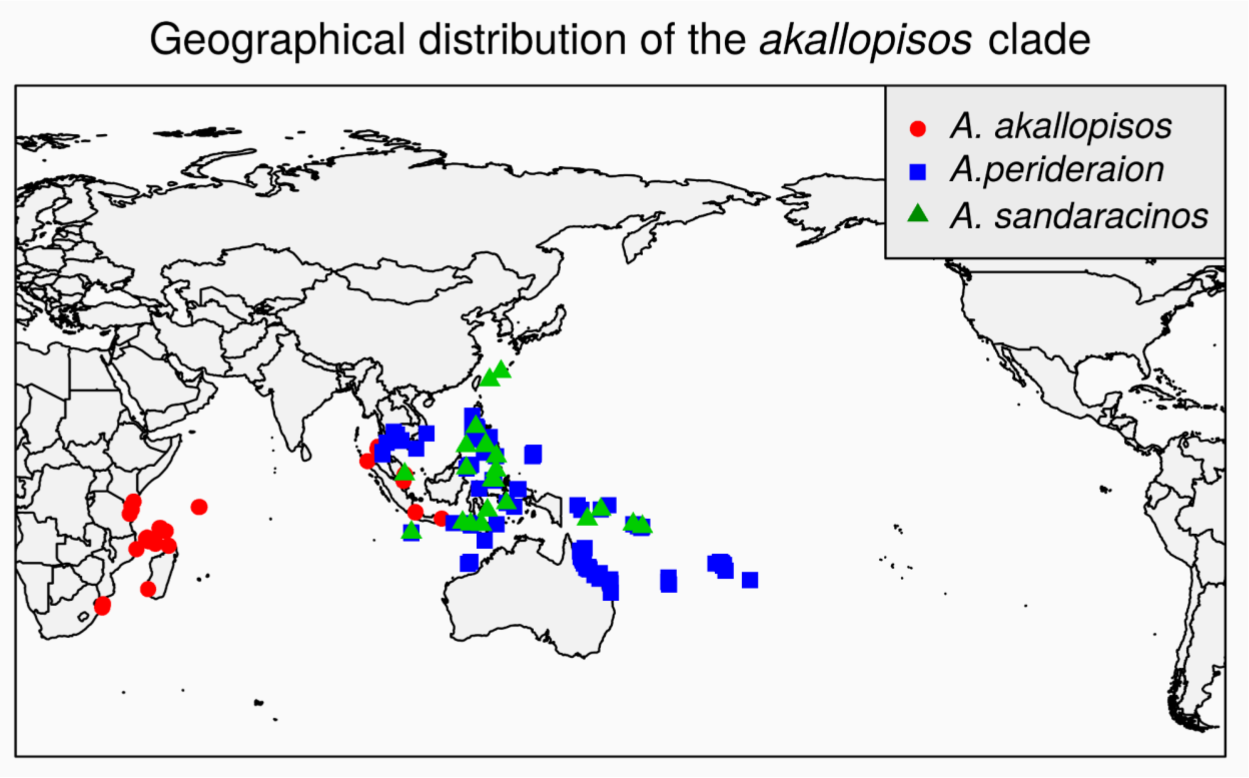

**Supplementary Figure S2:** SNPs density across the chromosomes.

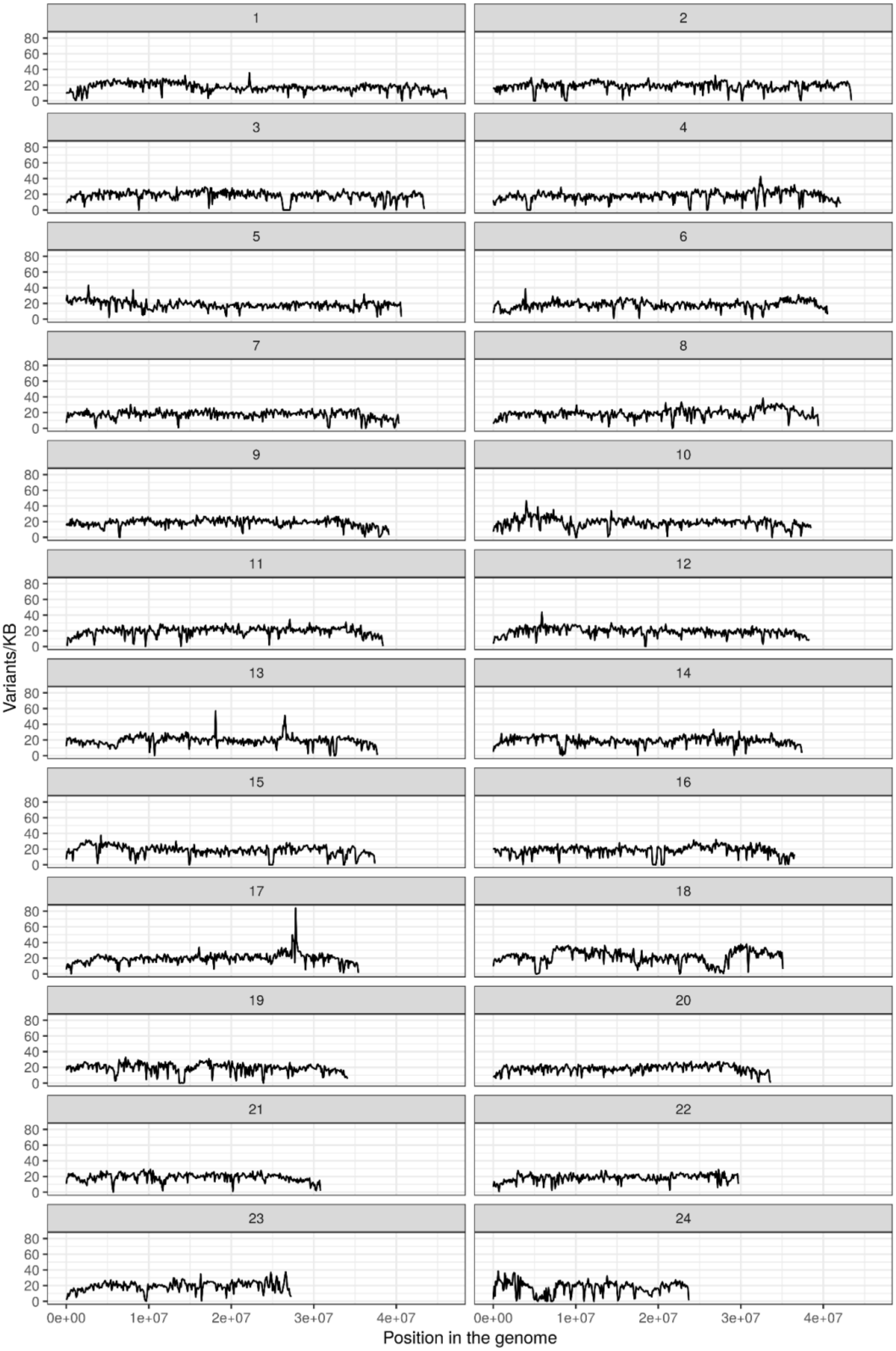

**Supplementary Figure S3. Admixture results for the individuals of the skunk complex obtained from PCAngsd.** In the admixture plot, individuals' ancestry proportions for two to five ancestral populations K are reported. Multiple runs with different seeds for each K were performed to ensure convergence. The highest likelihood solutions for each K are reported. Automatic selection of the best K by PCAngsd resulted in K=3.

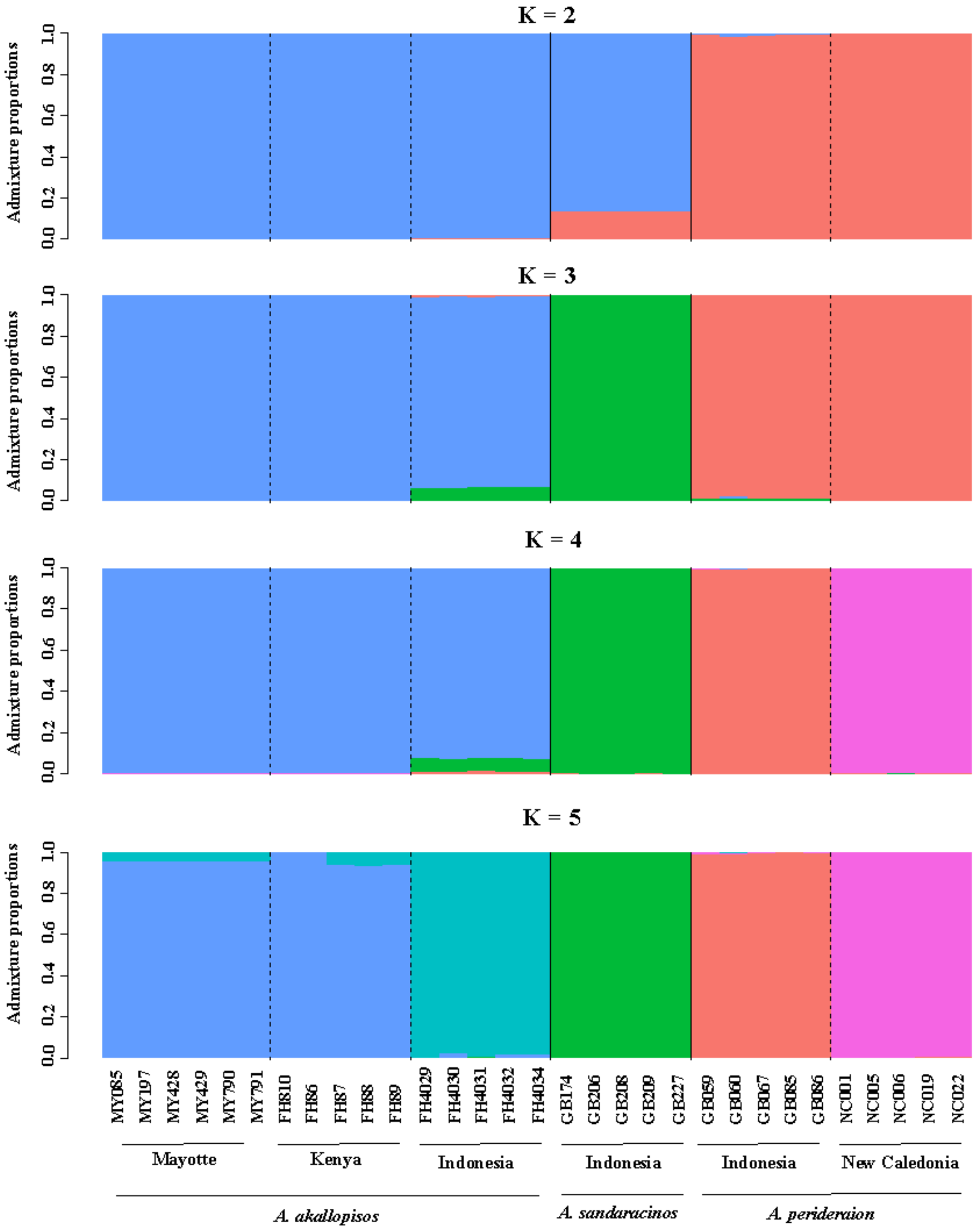

**Supplementary Figure S4. Absolute genetic divergence (*$d_{xy}$*) along the genome for all pairwise comparisons.** Red dotted lines represent the upper and lower 1% of the *$d_{xy}$* distribution. Aaka IND, Aaka KY and Aaka MY represent the populations of A. *akallopisos* from Indonesia, Kenya and Mayotte, respectively. Aper IND and Aper NC stand the populations of *A. perideraion* from Indonesia and New Caledonia, while Asand IND stands for *A. sandaracinos* from Indonesia. Values were calculated with the script *popgenWindows.py* (see Material and Methods).

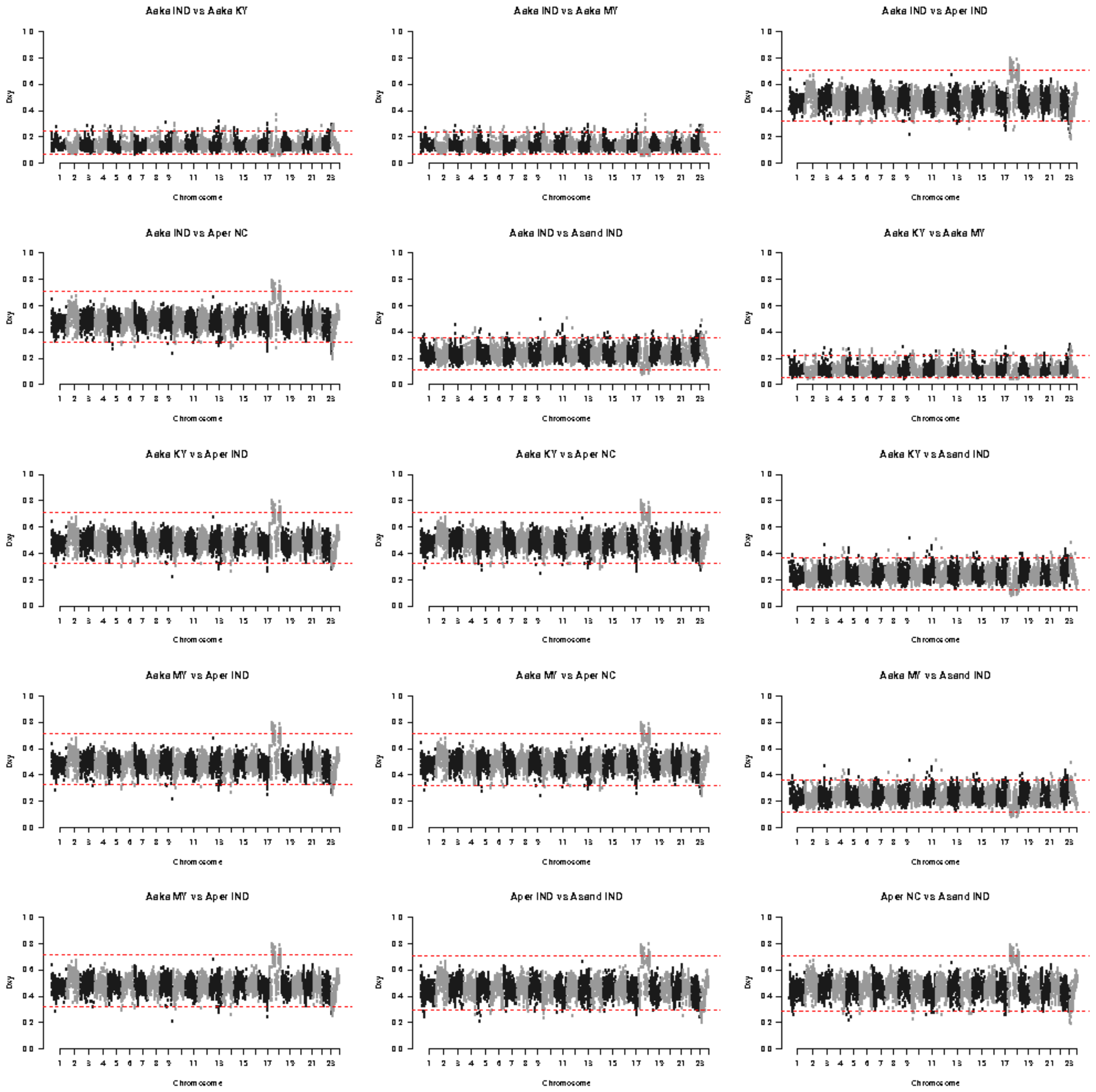

**Supplementary Figure S5. Absolute genetic divergence ($F_{st}$) along the genome for all pairwise comparisons.** Red dotted lines represent the upper and lower 1% of the $F_{st}$ distribution. Aaka IND, Aaka KY and Aaka MY represent the populations of A. *akallopisos* from Indonesia, Kenya and Mayotte, respectively. Aper IND and Aper NC stand the populations of *A. perideraion* from Indonesia and New Caledonia, while Asand IND stands for *A. sandaracinos* from Indonesia. Values were calculated with the script *popgenWindows.py* (see Material and Methods).

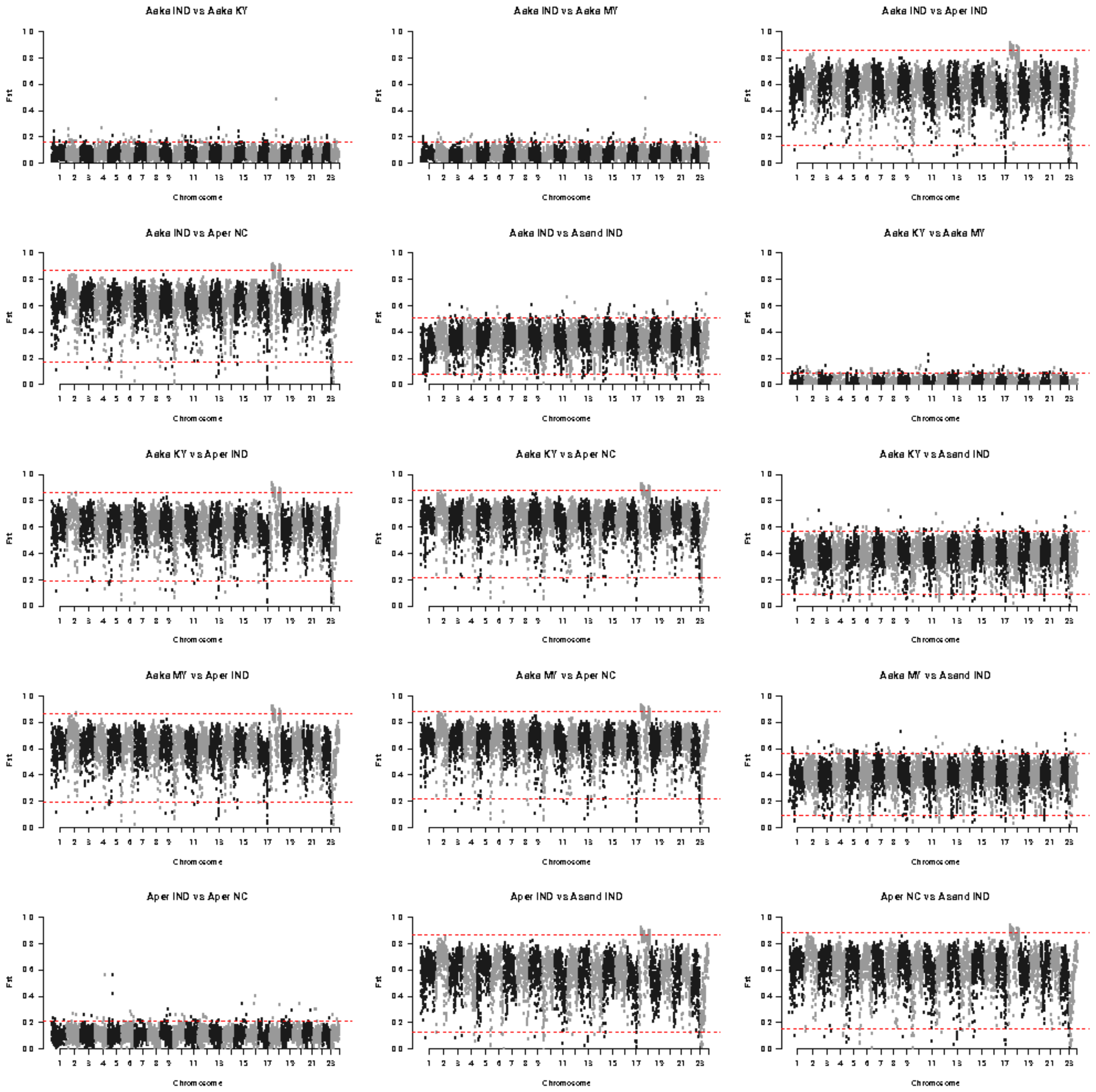

**Supplementary Figure S6. Nucleotide diversity (π) along the genome of all populations.** Red dotted lines represent the upper and lower 1% of the π distribution. Aaka IND, Aaka KY and Aaka MY represent the populations od *A. akallopisos* from Indonesia, Kenya and Mayotte, respectively. Aper IND and Aper NC stand the populations of *A. perideraion* from Indonesia and New Caledonia, while Asand IND stands for *A. sandaracinos* from Indonesia. Values were calculated with the script *popgenWindows.py* (see Material and Methods).

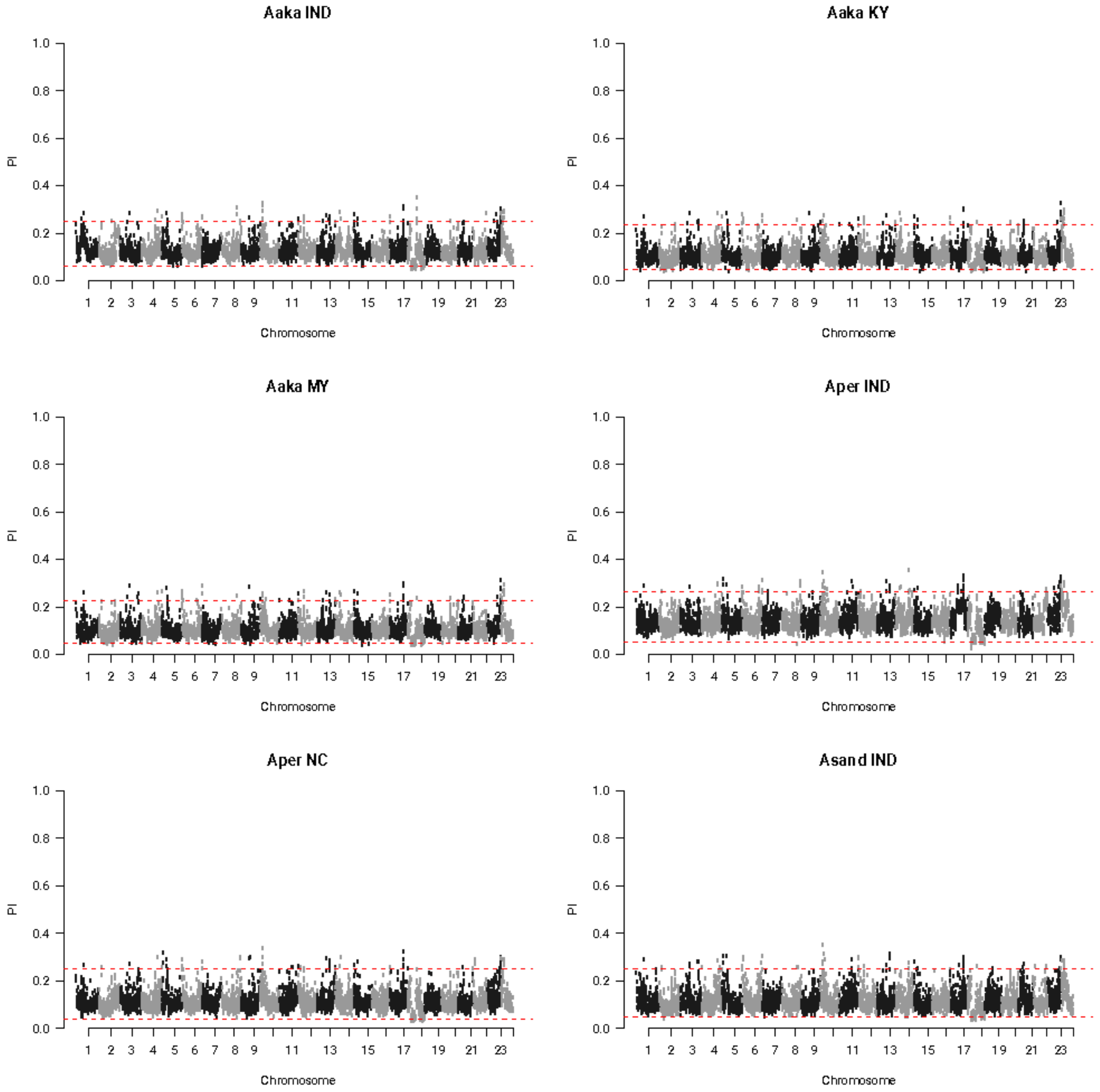

**Supplementary Figure S7.** ***Twisst*** **results along the chromosomes for the species-level analysis.** Colors correspond to the three possible topologies depicted in Fig. 2A.

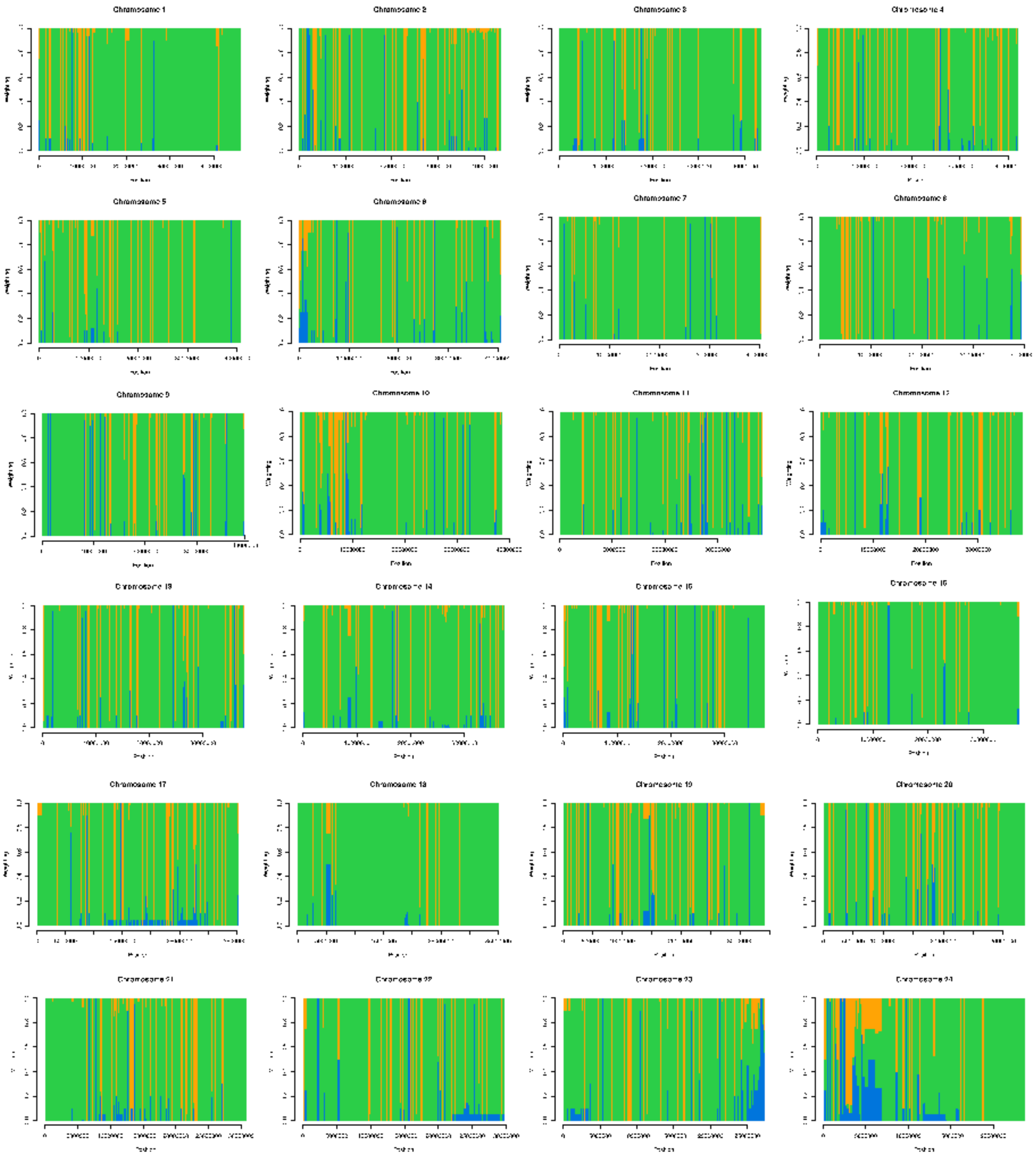

**Supplementary Figure S8. $f_d$ distribution along the chromosomes for the test of introgression between *A. perideraion* and *A. sandaracinos* individuals.** The red lines represent the 95th percentile of the $f_d$ distribution. Candidate region of introgression (CRI) are the regions in the upper 5% of the distribution and are represented in gray.

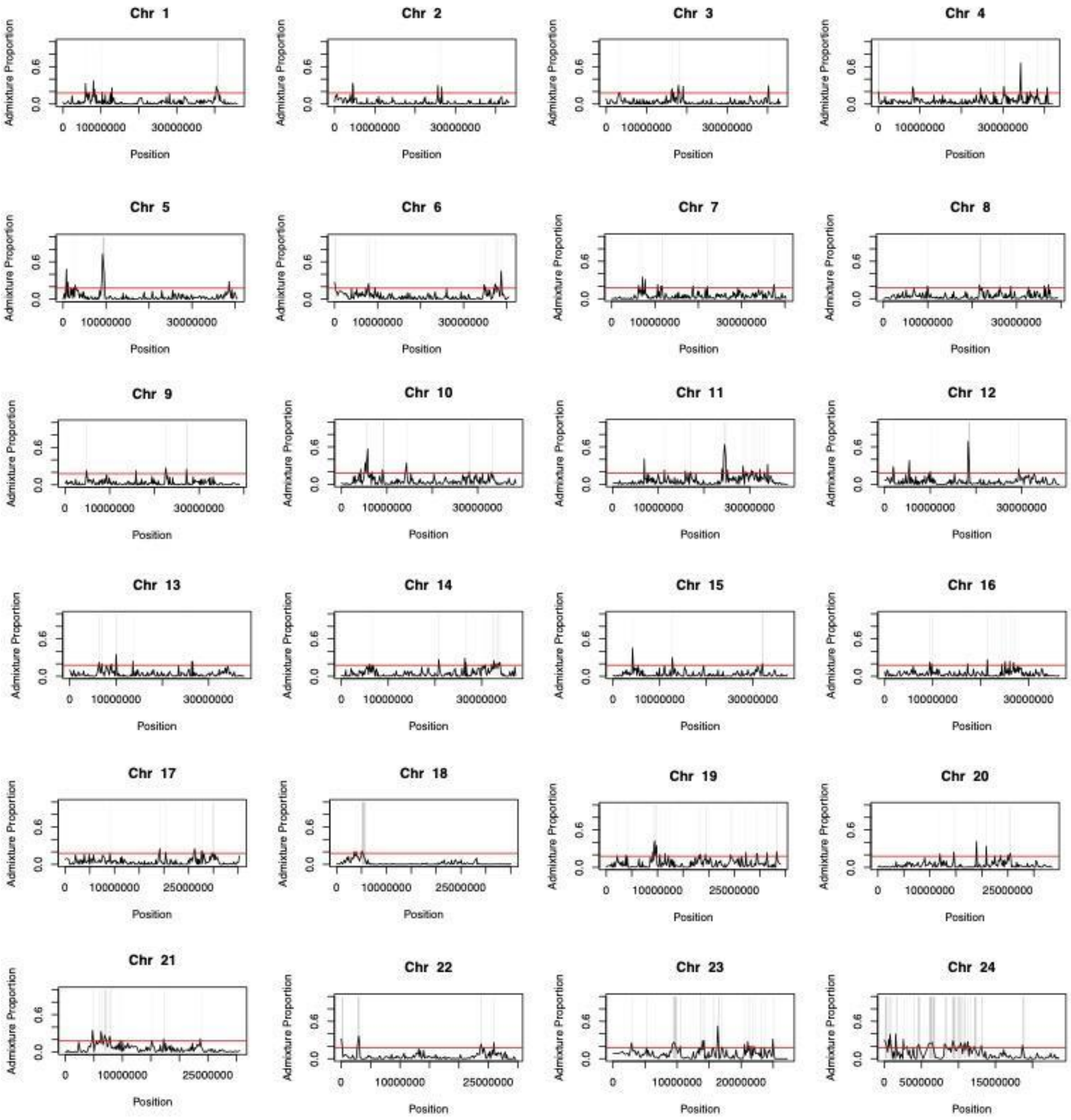
